# Quantification and discovery of sequence determinants of protein per mRNA amount in 29 human tissues

**DOI:** 10.1101/353763

**Authors:** Basak Eraslan, Dongxue Wang, Mirjana Gusic, Holger Prokisch, Björn Hallström, Mathias Uhlen, Anna Asplund, Frederik Ponten, Thomas Wieland, Thomas Hopf, Hannes Hahne, Bernhard Kuster, Julien Gagneur

## Abstract

Despite their importance in determining protein abundance, a comprehensive catalogue of sequence features controlling protein-to-mRNA (PTR) ratios and a quantification of their effects is still lacking. Here we quantified PTR ratios for 11,575 proteins across 29 human tissues using matched transcriptomes and proteomes. We analyzed the contribution of known sequence determinants of protein synthesis and degradation and 15 novel mRNA and protein sequence motifs that we found by association testing. While the dynamic range of PTR ratios spans more than 2 orders of magnitude, our integrative model predicts PTR ratios at a median precision of 3.2-fold. A reporter assay provided significant functional support for two novel UTR motifs and a proteome-wide competition-binding assay identified motif-specific bound proteins for one motif. Moreover, our direct comparison of protein to RNA levels led to a new metrics of codon optimality. Altogether, this study shows that a large fraction of PTR ratio variance across genes can be predicted from sequence and identified many new candidate post-transcriptional regulatory elements in the human genome.

## Introduction

Unraveling how gene regulation is encoded in genomes is central to delineate gene regulatory programs and to understand predispositions to diseases. Although transcript abundance is a major determinant of protein abundance, substantial deviations between mRNA and protein levels of gene expression exist (Liu *et al*, 2016). These deviations include a much larger dynamic range of protein abundances (García-Martínez *et al*, 2007; Lackner *et al*, 2007; Schwanhäusser *et al*, 2011; Wilhelm *et al*, 2014; Csárdi *et al*, 2015) and poor mRNA-protein correlations for important gene classes across cell types and tissues (Fortelny *et al*, 2017; Franks *et al*, 2017). Moreover, deviations between mRNA and protein abundances are emphasized in non-steady-state conditions driven by gene-specific protein synthesis and degradation rates (Peshkin *et al*, 2015; Jovanovic *et al*, 2016). Therefore, it is important to consider regulatory elements determining protein-per-mRNA copy numbers when studying the gene regulatory code.

Decades of single gene studies have revealed numerous sequence features determining protein-per-mRNA copy numbers affecting initiation, elongation, and termination of translation as well as protein degradation. Eukaryotic translation is canonically initiated after the ribosome, which is scanning the 5’UTR from the 5’cap, recognizes a start codon. Start codons and secondary structures in 5’UTR can interfere with ribosome scanning (Kozak, 1984; Kudla *et al*, 2009). Also, the sequence context of the start codon plays a major role in start codon recognition (Kozak, 1986). The translation elongation rate is determined by the rate of decoding each codon of the coding sequence (Sorensen *et al*, 1989). It is understood that the low abundance of some tRNAs leads to longer decoding time of their cognate codons (Varenne *et al*, 1984). However, estimates of codon decoding times in human cells and their overall importance for determining human protein levels are highly debated (Quax *et al*, 2015; Hanson & Coller, 2018). Secondary structure of the coding sequence and chemical properties of the nascent peptide chain can further modulate elongation rates (Qu *et al*, 2011; Artieri & Fraser, 2014; Sabi & Tuller, 2017; Dao Duc & Song, 2018). Translation termination is triggered by the recognition of the stop codon. The sequence context of the stop codon can modulate its recognition, whereby non-favorable sequences can lead to translational read-through (Poole *et al*, 1995; Tate *et al*, 1996; Bonetti *et al*, 1995; McCaughan *et al*, 1995). Furthermore, numerous RNA-binding proteins (RBPs) and microRNAs (miRNAs) can be recruited to mRNAs by binding to sequence-specific binding sites and can further regulate various steps of translation (Baek *et al*, 2008; Selbach *et al*, 2008; Guo *et al*, 2010; Hudson & Ortlund, 2014; Gerstberger *et al*, 2014; Cottrell *et al*, 2017). However, not only predicting the binding of miRNAs and RBPs from sequence is still difficult, but the role of most of these binding events in translation, if any, is poorly understood.

Complementary to translation, protein degradation also plays an important role in determining protein abundance. Degrons are protein degradation signals which can be acquired or are inherent to protein sequences (Geffen *et al*, 2016). The first discovered degron inherent to protein sequence was the N-terminal amino acid (Bachmair *et al*, 1986). However, the exact mechanism and its importance is still debated, with recent data in yeast indicating a more general role of hydrophobicity of the N-terminal region on protein stability (Kats *et al*, 2018). Further protein-encoded degrons include several linear and structural protein motifs (Ravid & Hochstrasser, 2008; Maurer *et al*, 2016; Geffen *et al*, 2016), or phosphorylated motifs that are recognized by ubiquitin ligases (Mészáros *et al*, 2017). Altogether, numerous mRNA and protein-encoded sequence features contribute to determining how many proteins per mRNA molecule cells produce. However, it is not known how comprehensive the catalogue of these sequence features is, nor it is known how they quantitatively contribute to protein-per-mRNA copy number.

To address these questions, Vogel and colleagues (Vogel *et al*, 2010) performed multivariate regression analysis to predict protein abundances from mRNA abundances and mRNA sequence features. According to their analysis, the major contributor to protein levels after mRNA abundance is the coding sequence, followed by the 3’UTR and then the 5’UTR. This seminal work was based on transcriptome and proteome data for a single cell type, Daoy medulloblastoma cells. Whether the conclusions can be generalized genome-wide and to other cell types remains an open question. Moreover, transcriptomics and proteomics technologies at the time were not as sensitive and quantitative as they are today. The mRNA abundances were estimated using microarray technology, which has a much poorer linear dynamic range than today’s RNA sequencing. Also, protein abundances were estimated using mass spectrometry-based proteomics, which was much less sensitive at the time, leaving reliable quantification only for 476 protein coding genes for further analysis. These 476 proteins were among the most abundant proteins, therefore leading to possibly strong analysis biases. Furthermore, this study was restricted to known sequence determinants of protein-per-mRNA abundances. No novel element was proposed.

Here we exploited matched proteome and transcriptome expression levels for 11,575 genes across 29 human tissues that we recently generated (see accompanying manuscript) to predict protein-to-mRNA ratios (PTR ratios) from sequence. We not only considered known post-transcriptional regulatory elements but also identified novel candidates in the 5’UTR, the coding sequence, and the 3’UTR, through systematic association testing. We also modeled the effect of codons on protein-to-mRNA ratio, leading to a new quantitative measure of codon optimality which we compared to existing measures. Our integrative model quantifies the contribution of all these elements on protein-to-mRNA ratio and predicts PTR ratios of individual genes at a relative median error of 3.2-fold. Finally, we provide initial experimental results to assess the functional relevance of these novel elements.

## Results

### Matched transcriptomic and proteomic analysis of 29 human tissues

We profiled by label-free quantitative proteomics and RNA-Seq adjacent cryo-sections of 29 histologically healthy tissue specimens collected by the Human Protein Atlas project (Fagerberg *et al*, 2014) that represented major human tissues (Fig 1A, see accompanying manuscript). We decided to model every gene with a single mRNA isoform because the number of genes with multiple quantified isoforms at the protein level was small (10% of the detected genes), and because of the inherent challenges and ambiguities of quantifying transcriptome and proteome data at the isoform level. For each gene, we defined the transcript isoform with the largest average protein abundance across tissues, measured as iBAQ values (Schwanhäusser *et al*, 2011), as its major transcript isoform. We used these major transcript isoforms for all tissues to compute all sequence features and to compute mRNA levels. The mRNA levels were estimated from RNA-seq data by subtracting length and sequencing-depth normalized intronic from exonic coverages. Subtracting intronic coverage led to slightly improved correlations between the mRNA and protein levels in every sample (Fig EV1) maybe because it better reflects the concentration of mature mRNAs, which are the ones exposed to the translation machinery. RNA-seq technical replicates were summarized using the median value. Requiring at least 10 sequencing-depth normalized reads per kilobase pair further improved the correlation between mRNA and proteins, likely because of the poorer sensitivity of proteomics for lowly expressed genes. We furthermore restricted the analysis to transcripts with a 5’ UTR and 3’ UTR longer than 6 nt so that all sequence features could be computed. Altogether, this analysis led to matched quantifications of protein and mRNA abundances for 11,575 genes across 29 tissues (Dataset EV1-4), where an average of 3,603 (31%) PTR ratios per tissue could not be quantified and were subsequently considered as missing values.

**Figure 1.**
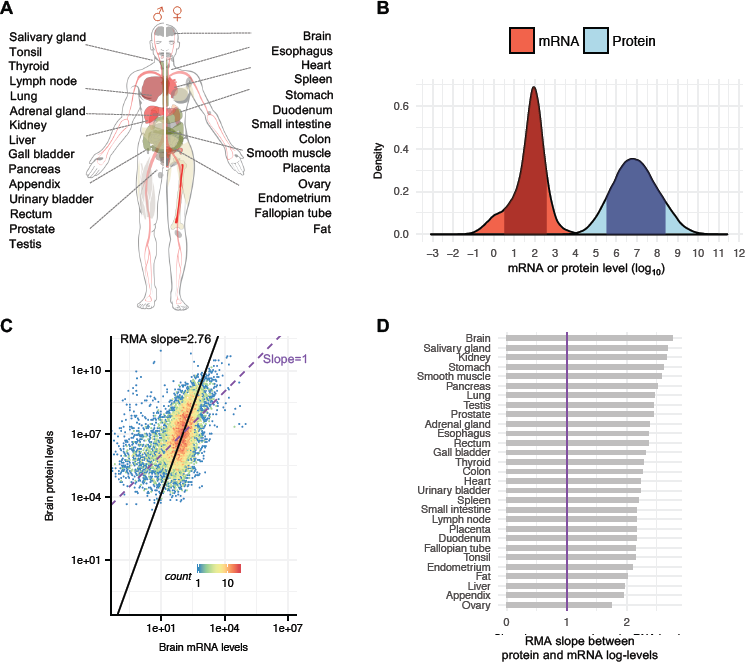
– Protein and mRNA levels across 29 human tissues. A The 29 tissues profiled in this study. B Distribution of log_10_ expression levels for mRNA (red) and protein (blue) across all genes and tissues and their 80% equi-tailed intervals (dark). C Protein levels (y-axis) versus mRNA levels (x-axis) in brain with ranged major axis (black) and a line of slope 1 passing through the point with median mRNA and protein levels (purple). D Ranged major axis slope for all 29 tissues in decreasing order. Vertical purple line marks the slope 1.

### Comparison of transcript and protein expression

Among these genes, the dynamic range of proteins spanned about 1 more order of magnitude than the dynamic range of mRNAs (125-fold for the 80% equi-tailed interval for the mRNA levels versus 794-fold for the proteins, Fig 1B). The enhanced dynamic range at the protein level implies that protein synthesis and protein stability play an important role in determining protein levels beyond mRNA levels. Moreover, we found a nearly quadratic relationship between mRNA levels and protein levels in every tissue (ranged major axis slope in a log-log plots of 2.76 in the brain, Fig 1C, and between 1.74 and 2.76 in all 29 tissues, Fig 1D, with Spearman correlations ranging from 0.42 to 0.62, Fig EV2, Materials and Methods). Hence, the protein copy number per mRNA is much larger for high abundant mRNAs than for low abundant mRNAs, consistent with observations in fission yeast (Lackner *et al*, 2007), baker’s yeast (García-Martínez *et al*, 2007; Csárdi *et al*, 2015), mouse (Schwanhäusser *et al*, 2011), and human (Wilhelm *et al*, 2014). This suggests that human genes encoding highly abundant proteins not only express mRNAs at high levels, but also encode post-transcriptional regulatory elements that favor high translation and protein stability (Vogel *et al*, 2010). We next exploited our systematic quantification of proteins and mRNA levels across 29 human tissues to estimate the contribution of known post-transcriptional regulatory elements on protein-to-mRNA ratio, within and across tissues, and to discover novel ones.

### Sequence determinants of protein-to-mRNA ratio

To identify and quantify sequence determinants of protein-to-mRNA ratio, we derived a model predicting tissue-specific PTR ratio from mRNA and protein sequence alone. The model is a multivariate linear model that includes a comprehensive set of mRNA-encoded and protein-encoded sequence features known to modulate translation initiation, elongation, and termination, and protein stability (Fig 2A-B, Materials and Methods, Dataset EV5). The model also includes sequence features that we identified *de novo* through a systematic testing of association between median PTR ratio across tissues and presence of k-mers, i.e. subsequences of a predefined length *k*, in the 5’ UTR, the coding sequence, the 3’ UTR, and the protein sequence (Materials and Methods). We report our findings below, going from 5’ to 3’. The effects of each sequence feature on PTR log-ratio are estimated using the joint model, thereby controlling for the additive contribution of all other sequence features (Dataset EV6).

**Figure 2.**
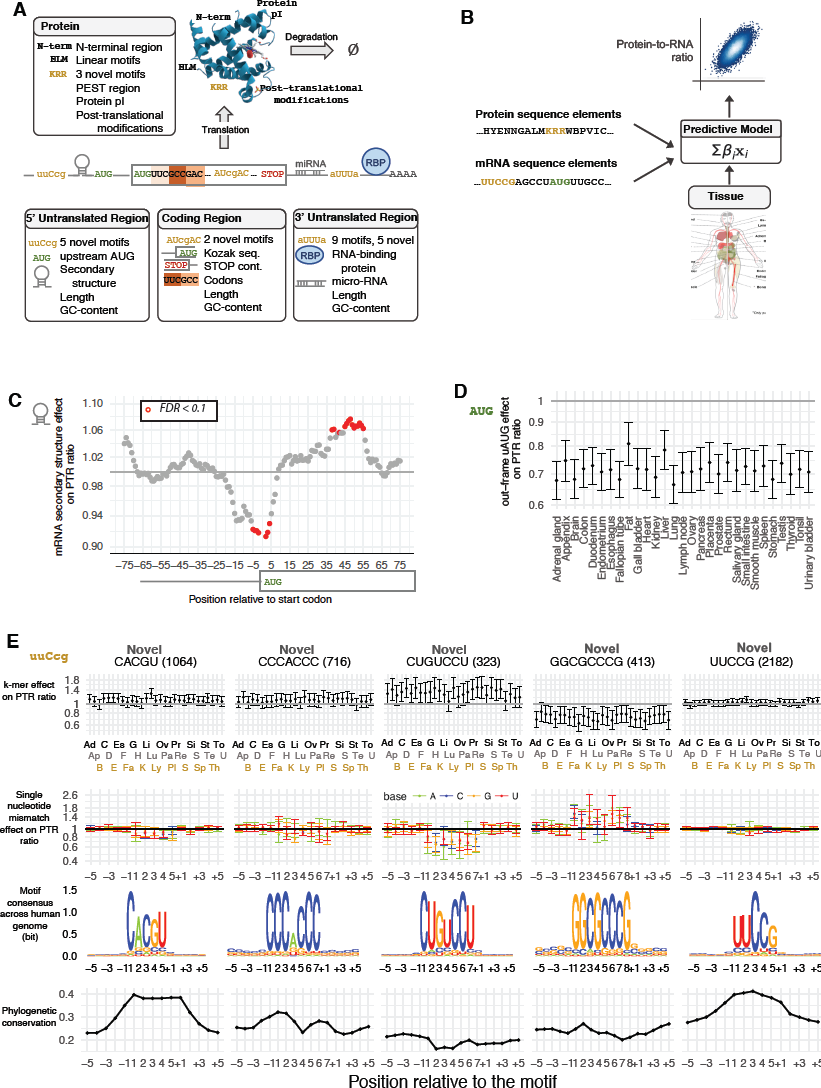
– Predicting PTR ratios from sequence and 5’ UTR results. A Sequence features of 5’ UTR, coding sequence, 3’ UTR, and protein sequence considered in the model. B The predictive model is a multivariate linear model that predicts tissue-specific PTR log-ratios using tissue-specific coefficients for the sequence features listed in (A) C Effect of log_2_ negative minimum folding energy of 51 nt window on median log_10_ PTR ratio across tissues corrected for all other sequence features listed in (A) (y-axis, Materials and Methods) versus position of the window center relative to the first nucleotide of the canonical start codon (x-axis) for genes with a 5’UTR and a coding sequence longer than 100 nt. Statistically significant effects at *P* < 0.05 according to Student t-test and corrected by Benjamini-Hochberg methods are marked in red. D Effect estimate (dot) and 95% confidence interval (bar) of the presence of at least one out-of-frame AUG in 5’ UTR on log_10_ PTR ratio corrected for all other sequence features listed in (A) (y-axis, Materials and Methods) per tissue (x-axis). E Each column corresponds to one of the 5 5’ UTR motifs (header with gene number in parentheses). From top to bottom. *First horizontal track*: Effect estimate (dot) and 95% confidence interval (bar) of the number of perfect match of the motif sequence on log_10_ PTR ratio corrected for all other sequence features listed in (A) (y-axis, Materials and Methods) per tissue (x-axis). *Second horizontal track*: Median effect (dot) and across tissue range (bar) of a single mismatch relative to the motif sequence on log_10_ PTR ratio corrected for all other sequence features listed in (A) (y-axis, Materials and Methods) per position around motif site(x-axis). Substitution with an A (green), C (blue), G (orange) and U (red). *Third horizontal track*: Position weight matrix logos showing information in bits (y-axis) computed across all motif instances in 5’ UTRs allowing for at most one mismatch. *Fourth horizontal track*: Average 100-vertebrate PhastCons score (y-axis, Materials and Methods) per position relative to the exact motif match instances in 5’ UTR (x-axis).

### mRNA 5’ UTR sequence features

Negative minimum RNA folding energy in 51 nt sliding windows, a computational proxy for RNA secondary structure, associated with a lower PTR ratio between 4 nt 5’ of the start codon and 4 nt 3’ of the start codon (Fig 2C, up to 9% decrease, FDR < 0.1, Materials and Methods). This inhibitory effect around the start codon is in agreement with mechanistic studies in *E. coli* showing that secondary structures around the start codon impair translation by sterically interfering with the recruitment of the large ribosome subunit (Kudla *et al*, 2009). In contrast, negative minimum folding energy in 51 nt windows associated positively with the PTR ratio between 38 nt and 55 nt 3’ of the start codon (Fig 2C, up to 8% increase, FDR < 0.1). This positive association is consistent with experiments showing that hairpins located downstream of the start codon facilitate start codon recognition of eukaryotic ribosomes in vitro (Kozak, 1990), supposedly by providing more time for the large ribosome subunit to be assembled.

Investigating every 3- to 8-mer in the 5’ UTR, while controlling for occurrence of other k-mers, revealed 6 k-mers significantly associated with PTR ratio at false discovery rate (FDR) less than 0.1 (Materials and Methods). These k-mers include AUG, the canonical start codon, for which at least one out-of-frame occurrence associated with about 18% to 32% lower median PTR ratios across tissues (Fig 2D), consistent with previous reports that out-of-frame AUGs in the 5’ UTR (uAUG) (Kozak, 1984) and upstream ORFs (uORF) (Morris, 2000; Calvo *et al*, 2009; Barbosa *et al*, 2013) associate with lower protein per mRNA amounts. No significant associations could be found for the 796 transcripts with only in-frame uAUGs (Fig EV3A). Among 2,483 transcripts with a single uAUG or uORF, a single out-of-frame uAUG is associated with a 22% reduced PTR ratio compared to a single out-of-frame uORF (Fig EV3B) possibly because ribosomes can re-initiate translation downstream with high efficiency after translating a uORF (Morris, 2000).

The other five 5’ UTR k-mers identified *de novo* had not been reported yet (Fig 2E). Occurrence of the 5-mer CACGU in the 5’ UTR associated with an increased PTR ratio of about 8%, with a maximum effect found in lung (25%). CACGU is present in the 5’ UTR of 1,064 (9%) of the 11,575 investigated genes, which are enriched for genes encoding proteins localizing to the nucleolus (Gene Ontology enrichment, Materials and Methods). Consistent with its hypothesized function in enhancing translation, the mismatch with any of the 5 nucleotides is rare and associates with a decreased PTR ratio by about 10% (Fig 2E). Moreover, instances of CACGU are more conserved than their flanking regions according to the PhastCons score (Siepel *et al*, 2005) computed across 100 vertebrates (Fig 2E, Materials and Methods). The second k-mer, CCCACCC, is present in 716 genes (6%) and associates with an increased PTR ratio of about 10% in the majority of the 29 investigated tissues. The two C triplets appear to be important because mismatches are rare in the genome and because these positions are more conserved than the central nucleotide and flanking nucleotides (Fig 2E). Minimum folding energy analysis displayed that CCCACCC favors to be in more unstructured regions of the mRNA (Materials and Methods, Fig EV4A), possibly because the binding trans-factor recognizes this motif within a specific mRNA secondary structure context as it has been shown for other RNA motifs (Aviv *et al*, 2006; Li *et al*, 2010; Luo *et al*, 2016; Taliaferro *et al*, 2016). The third k-mer, CUGUCCU, is associated with the strongest effect, associating with about a 33% increased PTR ratio in nearly every investigated tissue; mismatches for any of the 7 nucleotides is rare in the genome and associate with an entire loss of the effect. CUGUCCU is found in the 5’ UTR of 323 investigated genes (3%), enriched in genes of actin-mediated cell contraction pathway. The 8-mer GGCGCCCG, found in 413 genes (4%), associates with 25% lower PTR ratios in nearly every tissue and mismatches of any of the 8 nucleotides are rare and associate with a loss of the effect. GGCGCCCG prefers to reside in structured regions with relatively lower folding energy (Fig EV4B). The fifth k-mer, UUCCG is widespread (2,182 genes, 19%) and has a significant association with PTR ratio (about 9%, FDR < 0.1). Significance was lost when including RNA-binding protein data in the joint model, because of positive correlation with binding evidence for CSTF2, DDX3X, LARP4, and SUB1. Supportive for its functional role is the fact that mismatches to the consensus motif are rare for any nucleotide and that motif occurrences are conserved. UUCCG is enriched for genes of the RNA processing, translation, RNA binding pathways and for genes which are also components of the nucleolus and ribonucleoprotein complex.

### Start and stop codon context

Significant associations of individual nucleotides with PTR ratio were detected in [-6,6] nt around the start codon and in [-5,6] nt of the stop codon in at least one tissue (FDR<0.1, Fig 3A,B). At nearly every position of the start codon context, the nucleotide of the consensus sequence gccRccAUGG (Kozak, 1986) was the one associating with the largest effect, indicative of selection for efficient start codon recognition. The strongest effects were found at the third position 5’ of the start codon (25% lower PTR ratio for C than for the consensus A), recapitulating mutagenesis data (Kozak, 1986), and at the second nucleotide 3’ of the start codon (23% lower PTR ratio for A than for the consensus C). Moreover, effects of the start codon context on the PTR ratio were largely independent of the tissue (Fig 3A) consistent with a ubiquitous role of the start codon context likely due to structural interaction with the ribosome (Svidritskiy *et al*, 2014).

**Figure 3.**
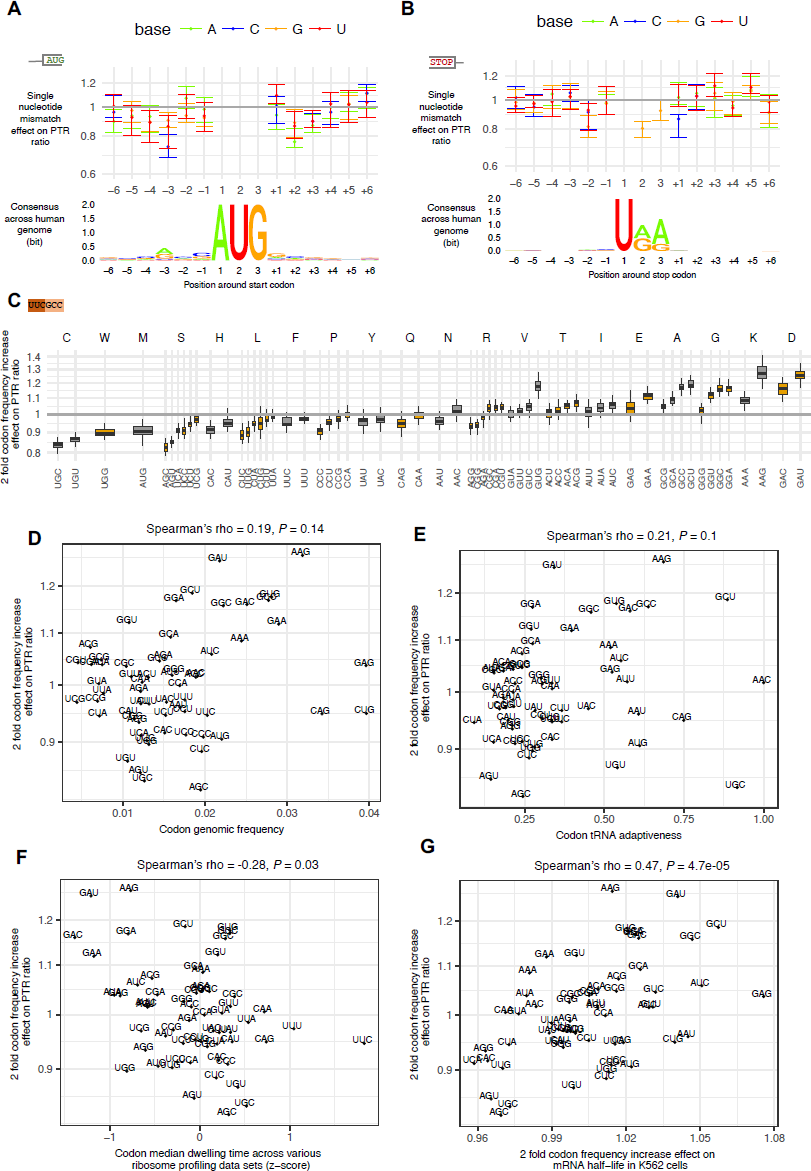
– mRNA coding region sequence features results. A Median effect (dot) and across tissue range (bar) of a single nucleotide mismatch relative to consensus sequence in a [-6, +6] nt window centered at first nucleotide of the canonical start codon.(*top*) Position weight matrix logo showing information in bits (y-axis) computed across all 11,575 transcripts. (bottom) B Same as A, centered at canonical stop codon. C Across tissue distribution of the estimated two-fold codon frequency increase effect on PTR ratio. Boxes represent 25%, 50% and 75% quantiles. D Codon genomic frequency (x-axis) versus effects of two-fold codon frequency increase on PTR ratio (y-axis). Spearman correlation is not significant. E Codon tRNA adaptiveness (x-axis) versus effects of two fold codon frequency increase on PTR ratio (y-axis). Spearman correlation is not significant. F Negative correlation between codon median dwelling time estimates across 17 ribosome profiling datasets (z-score) versus two-fold codon frequency increase on PTR ratio (y-axis). G High positive correlation between the effect of two-fold codon frequency increase on mRNA half-life in K562 cells (x-axis) versus PTR ratio (y-axis) across 29 tissues.

The opal stop codon UGA was significantly associated with the lowest median PTR ratio having in median 16% lower PTR ratios than the ochre stop codon UAA (Fig EV3C, *P* = 1.7×10^−5^). Moreover, the two most influential positions were the +1 nucleotide at which a C associates with 12% lower PTR ratios than the consensus G, and the −2 nucleotide, at which a G associates with 17% lower PTR ratios than the consensus A in median across tissues (Fig 3B). An inhibitory effect of a cytosine at the +1 nucleotide, which is observed for all three stop codons (Fig EV3D) is in line with previous studies in prokaryotes and eukaryotes (Poole *et al*, 1995; Tate *et al*, 1996; Bonetti *et al*, 1995; McCaughan *et al*, 1995) and structural data showing that a cytosine following the stop codon interferes with stop codon recognition (Brown *et al*, 2015), thereby leading to stop codon read-through. Moreover, our data indicate that the nucleotide at the −2 position, which is also reported to be highly biased in *E. coli* (Arkov *et al*, 1993), is significantly associated with PTR ratio and deviation from the consensus nucleotide A is associated with a reduced PTR ratio. Altogether, the start and stop codon contexts demonstrate the sensitivity of the PTR ratio analysis in detecting contributions to translation down to single nucleotide resolution.

### Codon usage

Among all investigated sequence features, codon frequency had the largest predictive power on PTR ratio in every tissue (between 14% and 21% of the variance, median 17%, Fig EV5A). The effects were large with a 2-fold increase in codon frequencies associating with 15% lower PTR ratios for the codons AGC and AGU and 29% higher PTR ratio for the codon GAU (Fig 3C). Part of this effect may be due to the amino acid content of the protein, which could influence protein stability, or due to amino acid content of the nascent polypeptide chain, which could influence translation elongation (Charneski & Hurst, 2013). However, alternative codons coding the same amino acid showed different effects on PTR ratios (Fig 3C) and the predictive power reduced when using amino acid frequency instead of codon frequency (from median 17% to median 15% of the variance, Fig EV5A). Moreover, although the codon frequency distribution at the 5’ end of the ORF is different from the rest of the CDS (Materials and Method, Fig EV5B) (Clarke & Clark, 2010; Tuller & Zur, 2015), the relative effects of codons in this region were highly correlating with the effects of the rest of the coding region (Materials and Methods, Spearman’s rho=0.38, P< 0.0024, Fig EV5C). These observations indicate that codon identity affects PTR ratio, possibly because cognate tRNA concentrations influence decoding time (Varenne *et al*, 1984). Moreover, effects of individual codons showed consistent amplitudes and directions across tissues (Fig 3C), rejecting the hypothesis of widespread tissue-specific post-transcriptional regulation due to varying tRNA pool content among different tissues (Plotkin *et al*, 2004; Dittmar *et al*, 2006).

Notably, codon effects on PTR ratio did not correlate well with previous codon optimality measures including frequency of codons in human coding sequences (Fig 3D, Spearman’s rho=0.19, P=0.14, Materials and Methods) and species-specific codon absolute adaptiveness ((Sabi & Tuller, 2017) Fig 3E, Spearman’s rho=0.21, P=0.1), which are based on genomic or transcriptomic data and strong modelling assumptions. We then asked whether codon effects on PTR ratios could be related to codon decoding times, whereby long decoding times would lead to lower translation output. Consistent with this hypothesis, codon effects on PTR ratios correlated significantly negatively with median codon dwelling time estimates across 17 ribosome profiling data sets (Fig 3F, Spearman’s rho=-0.28, *P*=0.03, Materials and Methods, Fig EV6) (Dana & Tuller, 2015; O’Connor *et al*, 2016). Moreover, we found that median codon effects on PTR ratios across tissues correlated significantly positively with codon effects on RNA stability in the K562 cell line (Spearman’s rho=0.47, *P*=4.7e-05, Materials and Methods, (Schwalb *et al*, 2016)). This agreement of effects between PTR ratios and mRNA stability is consistent with the fact that codon usage is causally affecting RNA degradation (Hoekema *et al*, 1987; Presnyak *et al*, 2015; Mishima & Tomari, 2016; Bazzini *et al*, 2016) in a way that is mediated by translation (Radhakrishnan & Green, 2016). Altogether, these results indicate that a PTR ratio-based measure of codon optimality is an attractive alternative to existing codon optimality measures and could help resolving some of the debates about the role of codon optimality in human cells.

Moreover, *de novo* motif searching in the 5’ and 3’ end of the coding sequence revealed two 6-mers significantly associated with PTR ratios, also when controlling for codon pair frequencies (two first principal components, Materials and Methods). The first 6-mer, AUCGAC, associates with on average 40% and 67% higher PTR ratios across tissues, when in frames 2 and 3, and in the first 300 nucleotides of the coding sequence (Fig EV7). The second 6-mer, GACUCU associates with between 50% and 30% lower PTR ratios across tissues, when in frame 2 and 3 in the last 300 nucleotides of the coding sequence (Fig EV7). How such motifs could function is unclear and there was no evidence for stronger conservation of these motif instances, irrespective of the frame (Fig EV7).

### Protein sequence features

Our model also includes protein sequence features. Although the N-terminal amino acid, known to affect protein stability via the N-end rule pathway, significantly associated with the PTR ratio (Fig EV8), the N-terminal amino acid was not significant in the joint model, likely because the effect was confounded with the start codon context. A recent study by Kats and colleagues (Kats *et al*, 2018) in yeast indicated that the mean hydrophobicity of the first 15 amino acids plays a more important role in protein stability than the N-end rule pathway. We observed that mean hydrophobicity of the first 15 amino acids significantly associated with the PTR ratios of 10 tissues (Fig 4) (5% higher PTR ratio on average, FDR <0.1), however positively, in apparent contradiction with its negative effect on protein stability in yeast (Kats *et al*, 2018). This may be due to the multiple roles of the coding region 5’ end in gene expression regulation, including translation (Tuller & Zur, 2015). We also considered protein surface charge-charge interactions because they can affect protein stability (Samantha S. Strickler *et al*, 2006; Chan *et al*, 2012), and because the charged polypeptides in the ribosome exit tunnel can influence ribosome elongation speed (Requião *et al*, 2017). Consistently, we observed that a one unit increase in the protein isoelectric point has a significant negative association with the PTR ratio (median 7%) in various tissues (Materials and Methods, Fig 4). Our analysis also confirmed, genome-wide, the negative effect on PTR ratios of PEST regions, degrons that are rich in proline (P), glutamic acid (E), serine (S), and threonine (T) (Rogers *et al*, 1986) that were present in 4,592 proteins (Materials and Methods), and quantified its median effect across tissue to a 14% lower PTR ratio (Fig 4).

**Figure 4.**
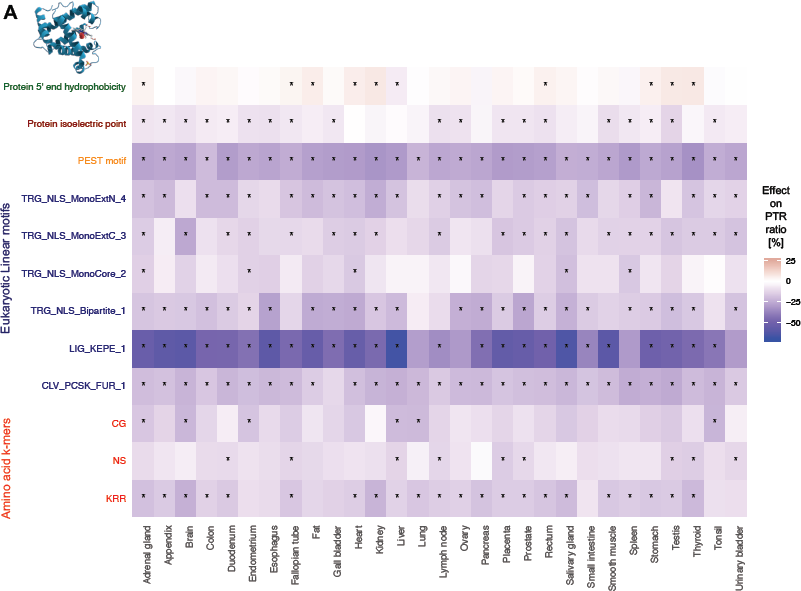
– Protein sequence features. Heatmap showing the tissue specific associations of protein sequence features with higher [0, + 25] (red gradient) or lower [-70, 0] (blue gradient) PTR ratios. Stars represent significance with FDR < 0.1.

De novo motif searching revealed two 2-mers and one 3-mer associating with lower PTR ratios (11%, 15%, and 8% median effects for CG, KRR, NS and respectively, Fig 4). The effect for KRR is consistent with association of stretches of positively charged amino acids directly upstream of high ribosome occupancy peaks in ribosome footprint data, suggesting that positively charged amino acids slow down translation (Charneski & Hurst, 2013). However, lysine (K), and arginine (R) are also the two amino acids recognized by cleavage sites of trypsin, the enzyme used to digest proteins prior to mass spectrometry. Although K and R as single amino acids do not stand out as negatively associated with the PTR ratio (Fig 3C), we cannot exclude a technical bias for the negative association of the 3-mer KRR with PTR ratios. Furthermore, we identified 6 linear protein motifs out of the 267 motifs of the ELM database (Dinkel *et al*, 2016) using a feature selection method (Materials and Methods). These 6 linear protein motifs contained all 4 nuclear localization signals of the ELM database which associated negatively with PTR ratios. It is unclear why nuclear localization signals could mechanistically lead to lower translation rates or higher protein degradation rates. These linear motifs are also KR-rich. Similar to the association of the 3-mer KRR, this could either reflect the effect of stretches of positively charged amino acid on slowing down translation or a technical bias due to the usage of trypsin as protein digestion enzyme (Materials and Methods, Fig4).

### mRNA 3’ UTR sequence features

*De novo* motif searching in the 3’ UTR revealed nine k-mers significantly associated with PTR ratios (FDR <0.1 Fig 5A and Fig EV9). This recovered 4 known mRNA motifs: the polyadenylation signal AAUAAA (Proudfoot, 1991), the AU-rich elements UAUUUAU (Cairrao *et al*, 2009) and AUUUUUA (Chung *et al*, 1996), and the binding site of the Pumilio family of proteins UGUAAAUA. The polyadenylation signal AAUAAA associated with between 5% and 27% increased PTR ratio across tissues (median 18%, <0.1 FDR, Fig 5A), was consistent with a role of polyadenylation signals in translation (Piqué *et al*, 2008). The evolutionary conserved AU-rich element UAUUUAU was found in 3,158 genes (27%) and associated with lower PTR ratios by about 9% consistently across tissues, in agreement with its function in mRNA destabilization and translational silencing (Chen & Shyu, 1995; Mukherjee *et al*, 2002). AAUAAA and UAUUUAU both favor to reside in unstructured mRNA sequence regions with higher folding energy (Fig EV10A,B). The Pumilio motif UGUAAAUA was found in 1,320 genes (11%), is highly conserved across species (Fig 5A) and is the binding target of members of the Pumilio family of proteins which regulate translation and mRNA stability in a wide variety of eukaryotic organisms (Parisi & Lin, 2000). Five novel motifs, GGCCCCUG, UACUAAGA, ACUCUG, CAACAGA, CGUGUGG were present in 571, 222, 3827, 381 and 380 genes out of the 11,575 inspected respectively (Fig 5A and Fig EV7). The effects of these motifs show little variation across tissues, and were on average, associated with a 13%, 47%, 9%, 7% higher PTR ratio except for CAACAGA which is associated with a 15% lower PTR ratio. CGUGUGG and GGCCCCUG occur in structured mRNA regions with lower folding energy (Fig EV10C,D), indicating a contextual preference of the factors recognizing these motifs. These motifs may or may not play a causal role in determining PTR ratios. Functional assays would be required to validate their specific roles.

**Figure 5.**
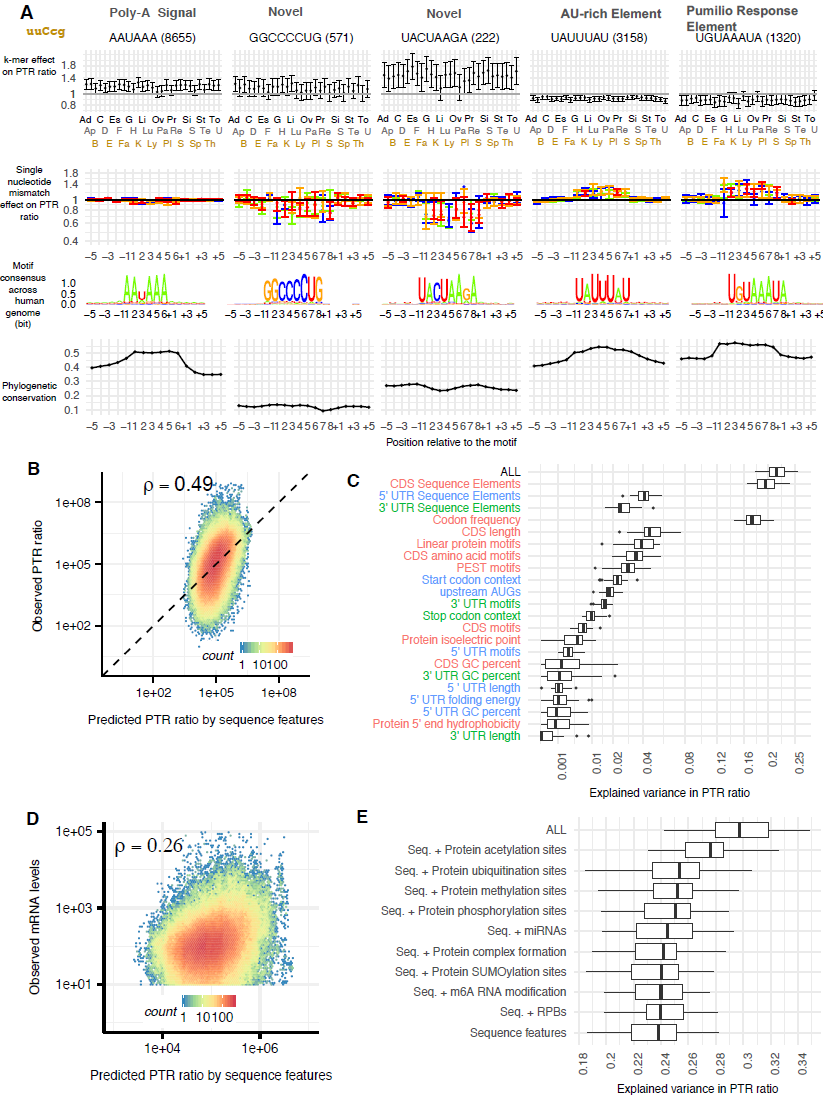
– 3’ UTR results and model summary. A Same as Figure 2E, for 5 3’ UTR motifs. B Measured PTR ratios for all genes in all tissues (y-axis) versus matching predicted PTR ratio by sequence features (x-axis). Colors stand for the density of genes. C Distribution of the explained variances (R^2^) in tissue specific PTR ratios only by each of the individual sequence features, as well as by the models combining them based on their mRNA region types (5’ UTR (blue), CDS (green) and 3’ UTR (green)) and altogether (ALL). Features are sorted based on their median explained variances. For improved visualization, the x-axis is shown with a squared root scale. D mRNA levels (y-axis) versus predicted PTR ratios by sequence features for all genes in all tissues. Colors stand for the density of genes. E Distribution of the explained variances (R^2^) in tissue-specific PTR ratios by the elastic net models which includes sequence features and features extracted from publicly available experimental data on protein post-translational modifications (acetylation, ubiquitination, methylation, phosphorylation, SUMOylation), mRNA m6A modification, protein complex membership, miRNA and RBP target sites.

### An interpretable model explaining PTR ratios from sequence

The multivariate linear model combining all these sequence features predicted PTR ratios at a median relative error of 3.2 fold on held-out data (10-fold cross-validation), which is small compared to the overall dynamic range of PTR ratios (200-fold for the 80% equi-tailed interval). This model explained 22% of the variance in median across tissues (min 18%, max 26%, Fig 5C). In comparison, a linear model based on codons alone explained 17% of the variance in the median across tissues (min 14%, max 21%, Fig 5C). This is followed by linear models based on protein sequence elements including the CDS length, the linear protein motifs, and our de-novo identified CDS amino acid motifs. The amount of explained variance is driven by the combination of effect size, frequency and variability of the feature across genes. Hence, sequence features which play a crucial role for translation, like the Kozak sequence can only explain 3% of the genome-wide PTR ratio variation, because it is already optimized for most of the genes in the genome. Also, the 5’ and 3’ UTR motifs explain a small fraction of the variance between genes although their effect size can be large (Fig 2E and Fig 5A), because they typically occur in a small number of genes. The predicted PTR ratios positively correlated with the mRNA levels (Fig 5D). Hence, our model supports the hypothesis that highly transcribed genes also have optimized sequences for post-transcriptional up-regulation, hence yielding higher amounts of proteins, consistent with earlier work by Vogel and colleagues (Vogel *et al*, 2010).

There are thousands of further sequence elements that could play a role in controlling the PTR ratios, including binding sites of any of the 2,599 catalogued human miRNAs (Chou *et al*, 2018), binding sites of the estimated about 1,500 RNA-binding proteins (Gerstberger *et al*, 2014), elements subject to mRNA modifications and post-translational modifications of certain amino acids. In this context, the derivation of a more comprehensive model of PTR ratio from sequence is difficult because i) the sequence determinants driving the binding of these factors and these modifications are poorly charted and ii) the number of regulatory elements is so large that one needs to rely on regularization techniques to prevent model overfitting, thereby losing model interpretability. To explore by how much the prediction of the PTR ratio from sequence could be improved in principle, we then considered a model that was not based on sequence alone but also included experimental characterization of such interactions and modifications of mRNA and proteins. This extended model included i) *N6*-methyladenosine (m6A) mRNA modification, an abundant modification enhancing translation (Wang *et al*, 2015), ii) binding evidence for 296 miRNAs from the mirTarBase database (Chou *et al*, 2018) with more than 200 targets in our dataset, iii) whether proteins are part of protein complexes, which is known to stabilize proteins (Mueller *et al*, 2015; Ishikawa *et al*, 2017), iv) binding evidence with 112 RNA-binding proteins (Van Nostrand *et al*, 2016), and v) phosphorylation, methylation, acetylation, SUMOylation and ubiquitination of certain amino acids (Hornbeck *et al*, 2015). Using a regularized model to cope with the large set of considered features (606 features, elastic net, Materials and Methods), this analysis showed that the explained variance can be increased to a median across tissues of 30% (Fig 5E min 24%, max 35%). Analysis of explained variance of the models based on individual feature groups indicate that post-translational modifications substantially contribute to PTR ratios (Fig 5E) and would constitute an important set of features to be modeled in more detail in the future.

### Independent confirmation of the model

As a starting point to assess the validity of our model and the derived predictions, we employed a three-fold approach, consisting of the confirmation of the prediction with an independent transcriptomic and proteomic dataset, a reporter assay measuring the dependence of the expression of a reporter protein to the presence of the 5’ or 3’ UTR sequence motifs, and a proteome-wide competition-binding assay to identify motif-specific RNA binding proteins and to measure their interaction strength. The effect of individual sequence features estimated on an independently generated dataset comprising matched RNA-Seq and proteomics data on 2,854 genes from 6 patient-derived fibroblasts (Kremer *et al*, 2017) agreed well with the median effects estimated across the 29 tissues (Materials and Methods, Spearman’s rho=0.64, *P*<2.2×10^−16^, Fig 6A). This was also true when restricting the analysis to the codon effects (rho=0.71, *P*<2.2×10^−16^), indicating that our new, regression-based measure of codon importance is reproducible across datasets.

**Figure 6.**
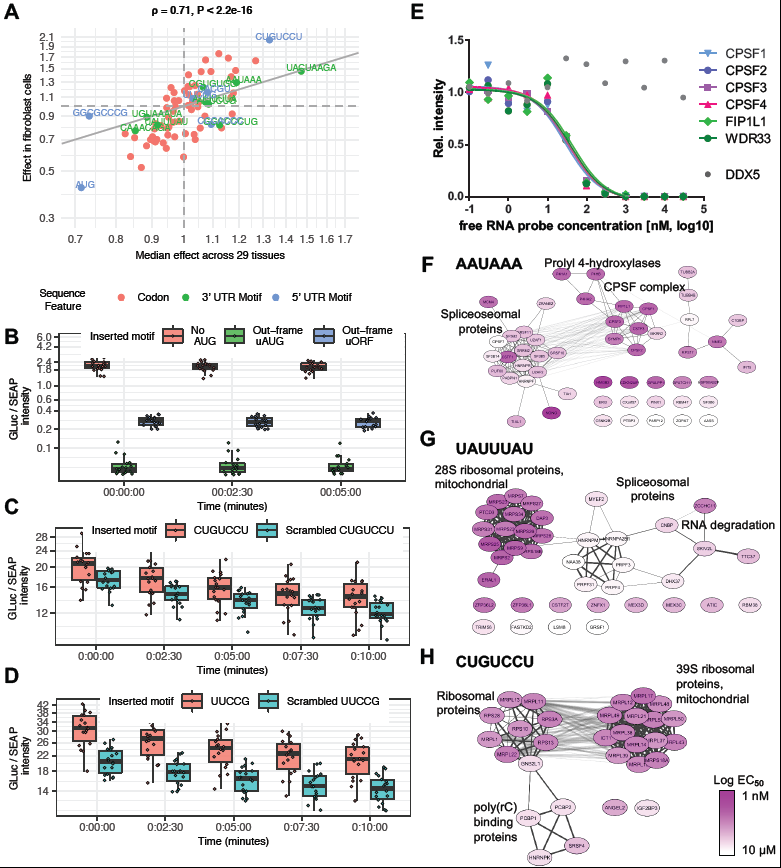
– Independent validations. A Comparison of codon (red), 3’ UTR (green) and 5’ UTR (blue) motif median effects on PTR ratio across 29 human tissues (x axis) and effects on PTR ratios in an independent matched proteome and transcriptome dataset Kremer et al. dataset (Kremer *et al*, 2017). B Reporter assay of the AUG in 5’UTR. Ratio of GLuc over SEAP intensities normalized per experiment (y-axis, Method) per time point (x-axis) and construct: no insertion (pink), inserted out-of-frame AUG (green) and inserted uORF, i.e inserted AUG with an inserted stop codon in frame in the 5’UTR (blue). Boxes mark the quartiles and whiskers 1.5 times the interquartile ranges. Original GLuc over SEAP intensities for this all all teste motifs in Fig EV11. C As in B for inserted 5’ UTR motif CUGUCCU (pink) or a scrambled version of it (blue, UUUGCCC). D As in B for inserted 5’ UTR motif UUCCG (pink) or a scrambled version of it (blue, CUUCG). E Proteome-wide competition-binding assay results for the polyadenylation signal motif AAUAAA and the Cleavage and polyadenylation specificity factor (CPSF) complex. F-H AAUAAA, UAUUUAU and CUGUCCU motif-specific RNA binding proteins (and their complex partners) and their interaction strength to the free RNA probe; node colour: pEC50; physical and functional interactions of proteins derived from STRING.

Next, we assessed the effects of motifs in a dual reporter assay in which the 9 tested motifs (Dataset EV7,8) were inserted in the 5’UTRs or 3’UTRs of Gaussia luciferase constructs (Materials and methods). The same plasmid also expressed a secreted alkaline phosphatase as control. This assay showed significant effects two positive controls: the out-of-frame upstream AUG and the out-of-frame upstream ORF, i.e. upstream AUG with an in-frame stop codon within the 5’UTR (Fig 6B, P < 0.0001). For the remaining tested motifs, control constructs containing scrambled versions of the tested motif were also assayed (Fig EV11, EV12, Dataset EV9). Two tested motifs (UUCCG and CUGUCCU) showed significant effects in the direction predicted by the model (Fig 6C,D, FDR<0.1, Materials and Methods). Most motifs have small predicted effects so that significance is difficult to attain in such assays. Taking this into account, four further motifs, including two positive controls, the AU-rich 3’ UTR motif UAUUUAU and the Pumilio response elements, as well the new motifs CCCACCC and GGCCCCUG, showed effects consistent with the model prediction (Fig EV11, EV12, Materials and Methods) in direction and amplitude.

Last, we investigated whether one of these motifs, the 5’ UTR motif CUGUCCU, was a potential recognition sites of RNA binding proteins. To this end, we performed a series of competition-binding assays with motif-containing RNA oligomers immobilized on Sepharose beads to capture RNA-binding proteins from HEK293 cell extracts in the presence of different concentrations of free motif-containing ligands. The captured RNA-binding proteins were analyzed by label-free quantitative proteomics. The resulting data allowed the estimation of EC50 values and dissociation constants (K_d_) akin to affinity chromatography-based chemical proteomics approaches (Médard *et al*, 2015; Bantscheff *et al*, 2007). The assay was also performed for two positive controls, the polyadenylation signal AAUAAA, and the AU-rich element UAUUUAU. Reversed or randomized sequences were used as negative controls. In total, we identified 253 proteins which directly or indirectly (e.g. as complex members) interact with the tested RNA motifs in a sequence-specific or sequence-independent manner. Of these, 88 proteins are annotated as RNA-binding proteins (RBPDB v. 1.3.1) and 247 proteins have annotations as RNA processing or binding proteins, nuclear localization, or (mitochondrial) ribosomal proteins (according to DAVID version 6.7 (Huang *et al*, 2009)). For further analyses, we defined sequence specific binding proteins based on their estimated K_d_ values: in order to be motif-specific, we required that the K_d_ of the protein-RNA interaction to be at least 10 times more potent than the next best motif or negative control. The competition binding experiments identified 50 motif-specific interactors of the consensus polyadenylation signal sequence AAUAAA (Table EV10), 32 of which are known to bind poly(A) tails and 12 have an annotated RNA binding domain. The results unambiguously recapitulate that the motif is bound tightly by the cleavage and polyadenylation specificity factor (CPSF) complex (Mandel *et al*, 2006)(Fig 6C). For the second positive control, the 3’ UTR AU-rich motif UAUUUAU (Chen & Shyu, 1995), the assay identified 38 interacting proteins. These include the zinc-finger RNA-binding proteins ZFP36L1 and ZFP36L2 (K_d_ of 24 nM, and 11 nM, respectively), which are known to destabilize several cytoplasmic AU-rich element (ARE)-containing mRNA transcripts by promoting their poly(A) tail removal or deadenylation, and hence provide a mechanism for attenuating protein synthesis (Hudson *et al*, 2004; Adachi *et al*, 2014). The competition assay also revealed interaction of many further proteins including ZCCHC11, MEX3C, MEX3D, CNBP, SKIV2, and TTC37 that are involved in mRNA decay consistent with the primary function of the AU-rich element.

The 5’ UTR motif CUGUCCU is one of the novel motifs with a predicted positive effect on the PTR ratio (1.33), which also showed as significant positive effect in the reporter assay. For this motif, we identified and quantified the interaction with 30 binding partners, including 19 proteins of the 39S mitochondrial ribosomal subunit, 5 proteins of the 40S ribosome complex, and 4 proteins with a KH domain known to be involved in splicing. The ribosomal proteins bind the 5’ UTR motif tightly with an affinity of 68 nM (+/- 31 nM std. dev.). One might speculate that the presence of this motif enhances the interaction of the 5’ UTR with the mitoribosome as well as with the small subunit of the cytoplasmic ribosome, which plays a key role in translation initiation (Aitken & Lorsch, 2012), leading to a higher efficiency in translation initiation. Two other proteins bound by the motif are ANGEL2, a protein known to bind the 3’ UTR of mRNAs resulting in their stabilization as well as IGF2BP3, a protein which may recruit and cage target transcripts to cytoplasmic protein-RNA complexes. Like other IGF2BPs, IGF2BP3 may thereby modulate the rate and location at which target transcripts encounter the translational apparatus and shield them from endonuclease attacks or microRNA-mediated degradation (Vikesaa *et al*, 2006; Wächter *et al*, 2013).

## Discussion

Our multivariate regression analysis quantified the contribution within and across tissues of 16 known sequence features representing 172 post-transcriptional regulatory elements, and further 15 novel motifs that we identified to be predictive of PTR ratios. Altogether the model predicts the PTR ratio of individual genes at a median precision of 3.2-fold from sequence alone, while the PTR ratio spans about 200-fold across 80% of the genes. For most known regulatory elements, the estimated effects were consistent with the literature, such as the effects of the secondary structures in the upstream CDS, upstream AUGs, individual nucleotides in the start and stop codon context, de-novo identified 3’ UTR motifs AATAAA, TATTTAT and TGTAAATA, providing support to the functional interpretability of the model. Moreover, this analysis led to the identification of novel candidate regulatory elements in 5’UTR and 3’UTR, whose effects are estimated to be in the range of well-known canonical motifs. Follow-up experiments provided initial functional support for these motifs.

To the best of our knowledge, the study by Vogel et al. 2010 (Vogel *et al*, 2010) is the only one that had integrated mRNA levels with an extensive list of known sequence features related to translation and protein degradation to predict human protein levels. In agreement with this study, we find that the coding sequence is the major contributor, followed by 3’ UTR and then 5’ UTR but we differ in the detailed features contributing to this observation. Compared to this seminal work, our study benefits from technological progresses made over the last decade. In particular, our proteomics data covers 11,575 proteins, whereas only 476 proteins could be used in the study by Vogel and colleagues. The larger dataset allowed us to model more sequence elements and to search for novel elements, some of which turned out to be present in a few hundreds of proteins.

Our regression approach led to a new codon metric, which quantifies the effect of individual codons on the PTR ratio, controlling for the additive effects of all other sequence features. Using this measure, codons are the lead explanatory variable explaining about 17% of the PTR ratio variance across genes. Codon usage plays an important role in translational efficiency due to varying translation elongation rates of different codons (Gustafsson *et al*,2012; Ingolia, 2014) and highly expressed genes contain relatively high proportions of codons recognized by abundant tRNAs with efficient codon-anticodon base-pairing. Based on this observation, several codon optimality metrics have been suggested (Sharp & Li, 1986; dos Reis *et al*, 2004; Pechmann & Frydman, 2013; Sabi & Tuller, 2017). However, all of these rely on some assumptions and simplifications, such as codon adaptation index defining a set of highly expressed genes as a reference set or tRNA adaptation index overlooking the supply and demand relationship for charged tRNAs. Recently, ribosome profiling experiments have enabled exploration of the codon-specific ribosome decoding rates (Ingolia, 2014) from ribosome footprints. Our acquired codon specific effects on the PTR ratio does not correlate with codon genomic frequency or tAI adaptiveness, while correlating well with the codon effects on ribosome decoding rates predicted by the RUST software (O’Connor *et al*, 2016) based on several ribosome profiling data sets. Furthermore, our PTR ratio-based codon metric also captures effects of amino acids on protein stability. Consequently, we suggest that the codon effect predicted from genome-wide PTR ratios, is a more reliable codon optimality metric compared to previous metrics. This metric could be named PTR-AI for protein-to-mRNA ratio adaptation index.

Our findings do not support the hypothesis of tissue type mediated codon optimality. It has been suggested that there is tissue-specific codon-mediated translational control due to differential synonymous codon usage in human tissue-specific genes, which correlates with varying tRNA expression among different tissues (Plotkin *et al*, 2004; Dittmar *et al*, 2006). On the contrary, also other studies found that their results do not support the evidence for optimization of translational efficiency by cell-type-specific codon usage in human tissues (Sémon *et al*, 2006; Rudolph *et al*, 2016). Our tissue specific codon effect predictions, which are acquired by fitting our model separately for each tissue, do not display high variation across tissues. This result is coherent with negligible tissue-specific enrichments of expressed codons in human transcriptomes (Sémon *et al*, 2006; Rudolph *et al*, 2016), showing that tissue-specific expression is neither due to the transcription nor to the translation of genes with particular codon contents. Further corroborating this finding, genes with high-effect codons tended to both have a high median level of protein expression and to be ubiquitously expressed. These genes were enriched for housekeeping functions. A possible explanation of these findings is that housekeeping genes evolve for optimal coding sequence to reach high protein expression levels. Because of the ubiquitous role of housekeeping genes, their codon content in turn constraints the pool of tRNA to be rather constant across tissues.

In every investigated tissue, protein-to-mRNA ratios were higher for genes with high mRNA expression levels yielding to an approximately quadratic relationship between protein and mRNA levels across genes. Our model explains in part this apparent amplification effect from sequence features, thereby showing that high protein expression levels are reached by the joint effect of high mRNA levels and genetically encoded elements favoring the synthesis and stability of proteins. A further fraction of this effect might be explained by regulatory elements that affect both the mRNA levels and protein per mRNA copy numbers. Codons are known to play such a dual role as they affect translation on the one hand, and mRNA stability on the other hand. The mechanistic basis for these cross-talks between translation and mRNA stability is not fully understood. Possibly, regression approaches as we employed here could help revealing further sequence elements acting on both levels. A similar super-linear relationship had been reported before for the unicellular eukaryotes baker’s yeast (Lackner *et al*, 2007) and fission yeast (Csárdi *et al*, 2015) yet appears to be absent in the prokaryote *E. coli*, for which mRNA and protein levels across genes obey a nearly linear relationship (Taniguchi *et al*, 2010). Prokaryotic transcription and translation are coupled processes, which do not allow post-transcriptional regulation to have an effective role in determining steady-state protein levels. In contrast, these two processes are highly uncoupled and have specialized mechanisms in eukaryotes, which is favored by the compartmentalization of eukaryotic cells. We suggest that the uncoupling of transcription and translation underlies a fundamental difference in the relationship between protein and mRNA levels across genes in eukaryotes compared to prokaryotes and may allow protein copy numbers of eukaryotic cells to span a much larger dynamic range. Further matched transcriptome and proteome datasets for a larger range of prokaryotes would help to support this model.

A comprehensive post-transcriptional regulatory code is important for interpreting regulatory genetic variations in personal genomes, and in genetic engineering for biotechnological or gene therapy applications. Our study provides an important contribution by modeling codon effects, identifying novel sequence elements with potential function and giving a framework to quantify and assess the role of new elements on protein-per-mRNA copy number. In the future, we expect further approaches including the analysis and integration of perturbation-based data and the mapping of post-translational regulatory elements, to complement and refine the present analysis.

## Materials and Methods

### Protein levels, mRNA levels, and PTR ratios

The protein data in MaxQuant file “proteinGroups.txt” is filtered such that the Reverse, Only.identified.by.site, and Potential.contaminant columns are not equal to \+”. Moreover, we restricted to unambiguously identified gene loci by requiring the number of ensembl gene IDs in the Fasta.headers column is equal to 1. We used as protein expression levels the per-tissue median centered log10 IBAQ values, settings missing values (NA) those IBAQ values equal to zero. To provide values in a consistent scale with the original values, the median of all log10 IBAQ values was further added.

About 10% of the genes were reported to have 2 or more transcript isoforms in the MaxQuant file “proteinGroups.txt”. We defined as major transcript isoform per gene, the transcript isoform reported in the MaxQuant file “proteinGroups.txt” that had the largest sum of IBAQ values across all tissues. We used these major transcript isoforms for all tissues, to compute all sequence features and to compute mRNA levels.

To estimate the mature mRNA levels, the exonic and intronic FPKM values were calculated separately and then the normalized intronic levels were subtracted from the normalized exonic levels. For exons, exon fragment counts per sample (i.e. each replicate of each tissue) are corrected by DeSeq2 library size factor and divided by the total exon length (in kilobase pairs). Likewise, the same is done for introns. The correction for intronic levels slightly improved the correlation between the mRNA and protein levels within samples compared to using exonic FPKMs maybe because it reflects better the cytosolic concentration of mRNAs. Finally, the technical replicates are summarized by taking the median value. We set a cut-off of 10 reads per kilo base pair for a transcript to be treated as transcribed, which further improved the correlation between mRNA and proteins, likely because of the poorer sensitivity of proteomics for lowly expressed genes. Tissue-specific PTR ratios were computed as the logarithm in base 10 of the ratio of the normalized protein levels over the normalized mRNA levels.

### Ranged major axis slope fitting

We fitted ranged major axis regression (Legendre *et al*, 2012), with protein level measurements (iBAQ (log_10_)) being the response variable and mRNA level measurements (normalized counts (log_10_)) being the single covariate, as implemented in the R package lmodel2.

### Sequence features

#### 5’ UTR folding energy analysis (secondary structure proxy)

The sequence spanning 100 nt 5’ and 100 nt 3’ of the first nucleotide of the canonical start codon were extracted for all transcripts with a valid PTR value in at least one tissue. The folding energies were computed via Vienna-RNAfold package (Lorenz *et al*, 2011) with 51 nt wide sliding window for each center position in [-75, +75] relative to the first nucleotide of the canonical start codon. The effect size and p-values of the log_2_-transformed negative minimum folding energy values at each position on median PTR across tissues was assessed individually with a linear regression model where all the analyzed sequence features were included as covariates. P-values were corrected for multiple testing using Benjamini Hochberg correction (Benjamini & Hochberg, 1995).

#### Kozak sequence and stop codon context analysis

Linear regression was performed on every nucleotide in a [-6,+6] nt window around the canonical start and stop codons.

### Codon usage

Codon usage was encoded as the log_2_ of the frequency of each 61 coding codons (number of codons divided by coding sequence length). Using the frequency in natural scale led to a decreased explained variance by 1%. In addition to that, codon pair frequencies were modeled in the design matrix as the first 2 principal components of the codon pair frequency matrix consisting of 3,721 features.

### Linear protein motifs

Linear protein motifs were downloaded from the ELM database (Dinkel *et al*, 2016) as regular expressions. We classified proteins as containing an ELM motif if the regular expression matched at least once in the protein sequence. Thereafter, we selected the ELM motifs significantly associating with PTR ratios in at least one tissue by utilizing LASSO feature selection (Tibshirani, 1996) where the PTR ratios were corrected for the core sequence features, which we defined as the motifs identified de novo, the 5’ UTR folding energies at positions 0 and +48, start codon context, codon frequencies, codon pair bias indicators, stop codon context, UTR and CDS region lengths, PEST motifs, protein isoelectric point and protein N-end hydrophobicity.

### N-terminal residue

The second residue of the protein sequence was extracted.

### Protein 5’ end hydrophobicity

Mean hydrophobicity value of the amino acids 2 −16 at the 5’ end of the protein were calculated by the hydropathy index per amino acid values reported in (Kyte & Doolittle, 1982).

### Protein isolectric point

Protein isolectric points for 11,575 protein considered in our model were computed with the IPC-isolectic point calculator software (Kozlowski, 2016).

### PEST-region

We classified protein sequences as “PEST-region containing” if the EMBOSS program *epestfind* (Rogers *et al*, 1986; Rice *et al*, 2000) identified at least one “PEST-no-potential” hit.

### De novo motif Identification

Similar to Eser et al (Eser *et al*, 2016) de novo motif identification was performed separately for 5’ UTR, CDS and 3’ UTR regions by using a linear mixed-effect model in which the effect of each individual k-mer on the median PTR ratios across tissues was assessed while controlling for the effect of the other k-mers (random effects), region length and region GC percent (fixed effects). The model was fitted with the GEMMA software (Zhou *et al*, 2013). Motif search was executed for k-mers ranging from 3 to 8 and the p-values were adjusted for multiple testing with Benjamini-Hochberg’s False Discovery Rate. Significant motifs at FDR < 0.1 were subsequently manually assembled based on partial overlap.

### Multivariate linear model (interpretable model)

The multivariate linear model for tissue *j* is y_ij_ = β_0j_ + X_i_β_j_ + є_ij_ where X is a matrix of sequence feature predictors which contains 61 features for individual codon frequencies (in log2 scale), 36 features for kozak sequence position-nucleotide pairs, 38 features for stop-codon context position-nucleotide pairs, 3 features for CDS, 5’ UTR and 3’ UTR lengths (in log2 scale), 3 features for CDS, 5’ UTR and 3’ UTR GC percentages, 9 features for 3’ UTR motifs, 6 features for 5’ UTR motifs (including upstream AUG), 2 features for CDS motifs, 3 features for CDS amino-acid motifs, 6 features for linear protein motifs, 3 features for 5’ UTR folding energy, 2 features for codon pair bias, 1 feature for pest motifs, 1 feature for protein isoelectric point, 1 feature for protein n-terminal hydrophobicity.

To predict the tissue-independent effects of the sequence features, we utilized a partial-pooling scheme with a multilevel varying-intercept multivariate regression model y_ij_ = β_0[j]_ + X_i_β + є_ij_ where the intercepts varied by tissue while the slopes of the sequence features kept equal across tissues.

The explained variance (R^2^) of the PTR ratio by the sequence features was obtained by 10-fold cross validation where in each fold the held-out data were used to have the PTR ratio predictions based on the linear regression model fit obtained from the remaining 9 partitions.

## Motif analysis

### Tissue specific motif effects

In the design matrix all of the de novo identified motifs except “AUG” and “AAUAAA” are encoded as the number of motif sites in the sequence of the mRNA region (i.e 5’ UTR, CDS, 3’ UTR). “AUG” and “AAUAAA” are encoded as binary, that is whether the motif is available in 5’ UTR and 3’ UTR regions respectively. The tissue-specific effect of the motif is assessed by fitting all sequence features considered jointly in the linear model, with the tissue specific PTR ratios as being the response variables.

### Gene ontology enrichment

Enrichment for gene ontology categories (Ashburner *et al*, 2000) as of January 21, 2016 was performed using the Fisher exact test and corrected for multiple testing using the Benjamini Hochberg correction (Benjamini & Hochberg, 1995).

### Motif 1 nucleotide mismatch effects

In order to assess the effect of the 1 nucleotide mismatch effect on PTR ratios, all motif occurrences with at most 1 mismatch, irrespective of the position, are found. The effects of 1 nucleotide mismatches with respect to consensus sequence are estimated with a linear model where the response is the residuals of tissue-specific PTR ratios corrected for other sequence features considered in the joint model and the design matrix contains the total number of motifs with at most 1 nucleotide mismatch as well as the number of mismatch instances per position-nucleotide type.

### Motif 1 nucleotide mismatch logos

The sequences in a [-5,+5] nucleotides window around each motif instance with at most 1 nucleotide mismatch were obtained from transcript mRNA sequences. The logos were created with R ggseqlogo package.

### Motif conservation analysis

Phylogenetics conservation scores for human annotation hg38 (phastConst100way from http://hgdownload.cse.ucsc.edu/goldenpath/hg38/phastCons100way), which reports conservation across 99 vertebrates aligned to the human genome, was downloaded and the conservation scores per nucleotide were extracted for each of the motif instances without any mismatch.

### Motif position density analysis

After filtering out the UTR regions which are shorter than 200 nucleotides, motif density per 20 nucleotide window was calculated by using the number of motif instances in a window normalized by theoretical possible k-mer numbers at each position.

### Motif region folding energy analysis

For each motif site, the [-250, +250] nt window sequences centered at the motif are extracted. Across each of these 500 nucleotide sequences, the folding energies of 51 nucleotide long sequences were computed by Vienna-RNAfold package (Lorenz *et al*, 2011) in a 1 nucleotide sliding windows approach. Minimum energy values are transformed to negative absolute log_2_ scale.

### Codon usage analysis

Codon frequency in human coding sequences were obtained from https://www.genscript.com/tools/codon-frequency-table. Codon decoding times for 16 human ribosome profiling datasets were obtained from RUST values (O’Connor *et al*, 2016). We estimated decoding time using the RUST ratio defined by the RUST A-site values over the RUST expected value (personal communication with Patrick O’Connor). We also included decoding times in the HEK293 cell line estimated by (Dana & Tuller, 2015). To estimate codon effects on mRNA half-life, we first called a major isoform as the highest expressed isoforms of Gencode v24 coding transcripts in the total RNA samples of Schwalb and colleagues (Schwalb *et al*, 2016) according to Kallisto (Bray *et al*, 2016). The half-life was estimated as the ratio of 5 min labelled TT-seq sample over total RNA seq sample (2 replicates) after correcting library size with spike-in. We then fitted a linear model with log_2_ RNA half-life as response variable against log_2_ frequency of codons with region length and GC content of 5’ UTR, CDS and 3 ‘UTR as further covariates.

### Coding sequence 5’ end codon frequency analysis

Considering 10,778 transcripts with CDS length greater than 460 nucleotides, starting from the second codon the log_2_ frequencies of 61 coding codons are calculated in each of the 11 non-overlapping 15 codon long windows. For Fig EV5B, the frequency values are centered per codon across windows.

In order to compare the effect of 2-fold codon frequency increase in the first window (codons from 2 to 16) versus the effect of the 2-fold codon frequency increase in the rest of the coding sequence, the codon frequencies of the whole coding sequence is replaced by the respective frequency values in the global model. The median effect of the codons across tissues are displayed in Fig EV5C.

## Non-sequence features

### m6A mRNA modification

We classified mRNAs as m6A modified if at least one m6A peak for the same gene locus in untreated HepG2 cell line was reported in the Supplementary Table 6 of Dominissini et al. (Dominissini *et al*, 2012).

### Protein complex membership

We classified each protein as a protein complex member if it was a subunit of at least one annotated protein complex in the CORUM (Ruepp *et al*, 2010) mammalian protein complex database (release version 02.07.2017).

### Protein post-translational modification

We downloaded protein acetylation, methylation, phosphorylation, SUMOylation and ubiquitination data from the Phosphosite database (release version 02.05.2018) (Hornbeck *et al*, 2015) and calculated the number of modification site per modification type for each protein. For proteins whose modification information was not available in the downloaded data set, we assigned 0 instead. The covariate for each of these features was defined as the log_2_ of the number of modifications plus 1 (pseudocount). ***RNA-binding protein targets:*** We classified transcripts as targets of 112 RBPs if they contained at least one peak in the eCLIP data set of Van et al. (Van Nostrand *et al*, 2016) as processed earlier (Avsec *et al*, 2018).

### miRNA targets

Many miRNAs of miRTarBase database (Chou *et al*, 2018) have very few reported targets, leading to no improved explained variance. We therefore filtered for the miRNAs which have at least 200 experimentally validated target genes in our data set and classified the mRNAs accordingly as targets for these miRNAs.

### Independent matched transcriptome-proteome dataset

We used data from Kremer et al. (Kremer *et al*, 2017). As originally reported, these data showed strong technical effects. To be on the safe side, we restricted the analysis to 6 samples (sample IDs: #65126, #73804, #78661, #80248, #80254, and #81273) that belonged to same cluster (Kremer *et al*, 2017).

### Elastic net model (regularized model)

For predicting the PTR ratios with the extended (sequence features and experimentally characterized features) feature set, we utilized elastic net models (Zou & Hastie, 2005) by the use of the R glmnet package (Friedman *et al*, 2010). We performed model selection on the training data (randomly selected 66% of all transcripts) where we set the parameter α to be 0.5 in 10-fold cross-validation. For the prediction of the test response (remaining 33% of all transcripts), we selected the model with the largest  value whose mean squared error (MSE) is within 1 standard error of the minimum.

### Validation of RNA motifs using a GLuc/SEAP reporter assay

We assayed the expression of one reporter gene on a plasmid (Gaussia luciferase, GLuc) as a function of the presence of a motif, while the second, constitutively expressed reporter gene (secreted alkaline phosphatase, SEAP) was used as internal control for variation in transfection efficiency and plasmid number. The pEZX-GA01 vector and the pEZX-GA02 vector (GeneCopoeia), containing Gaussia luciferase (GLuc) as a reporter and a constitutively expressed secreted alkaline phosphatase (SEAP) as an internal control, served as basic vectors of our 5´UTR and 3´UTR constructs respectively. We cloned the SV40 promoter and the 5´UTR motifs upstream of Gaussia luciferase ORF between the EcoRI and the XhoI sites of the pEZX-GA01 vector and 3´UTR downstream of the Gaussia luciferase stop codon between the EcoRI and the XhoI sites of the plasmid pEZX-GA02. The list of the motifs and controls used in the study is available in Dataset EV7. The luciferase assay was performed using Secrete-Pair™ Dual Luminescence Assay Kit (GeneCopoeia). A total of 100,000 HEK293 FT cells per construct were plated in 12-well plates. The following day, cells were transfected with 1 μg of DNA of each construct using Lipofectamine 2000 transfection reagent (Life Technologies) according to manufacturer´s protocol. The medium was changed 24 h after transfection and the cell culture medium was collected 48 h after transfection. GLuc and SEAP activities were measured with the Secrete-Pair™ Dual Luminescence Assay Kit (GeneCopoeia) according to the manufacturer´s protocol on Cytation3 imaging reader (BioTek). Each construct was measured in three technical and three biological replicate in two independent experiments with intensity measurements collected on 5 time points (0, 2, 4, 7, and 10 minutes).

For each motif separately, we assessed the per experiment significance of the effect of the motif versus its scrambled counterpart with a two-level nested ANOVA model fitted as a mixed effects model in which the response is the log-intensity ratio of GLuc over SEAP. In each of these motif specific models, the motif type (motif versus scrambled motif) is treated as the fixed effect while the replicate identifier is treated as the random effect (Fig EV11, EV12). Thereafter, we combined the P-values of the replicate experiments using Fisher’s method (Fisher, 1925) and corrected for multiple testing with Benjamini Hochberg correction (Benjamini & Hochberg, 1995).

### Competition-binding assay to identify RNA motif-binding proteins

The experiments were performed in three biological replicates and in two independent experiments. HEK293FT cells were grown in DMEM medium supplemented with 10% (v/v) FBS, 1% (w/v) non-essential amino acids, 1% (w/v) L-glutamine and 1% (w/v) G418 (geneticin, Thermo). Confluent cells were harvested by mechanical detachment followed by centrifugation and washing with cold Dulbecco’s phosphate buffer saline containing Ca^2+^ and Mg^2+^. Cell extraction and preparation of the lysate for the competition-binding assay was performed as described (Médard *et al*, 2015). The preparation of RNA-beads for affinity purification of RNA-binding proteins was performed as follows: NHS-Sepharose beads (Amersham Biosciences) were washed with DMSO(4 × 10 mL/mL beads) and reacted with RNA oligos with 5’ amino modified C6 (Dataset EV10), 50 nmol/mL beads; Integrated DNA Technologies, Inc.) for 20 h on an end-over-end shaker in the dark in the presence of triethylamine (30 μL/mL beads) in DMSO and H_2_O (1.8 vol of DMSO and 0.2 vol ddH_2_O for 1 vol of beads). Next, aminoethanol (50 μL/mL beads) was added and the mixture was kept shaking for an extra 20 h in the dark. The beads were washed with DMSO (10 mL/mL beads) and ethanol (3 × 10 mL/mL beads) and stored in ethanol (1 mL/mL beads) at 4 °C.

The competition binding assay itself, quantitative label-free LC-MS/MS analysis as well as curve fitting was also developed according to (Médard *et al*, 2015). Briefly, the diluted cell lysates (2.5 mg of total proteins/well) were incubated for 1h at 4 °C in an end-over-end shaker with 0 nM (water control), 0.3 nM, 1 nM, 3 nM, 10 nM, 30 nM, 100 nM, 300 nM, 1 μM, 3 μM, 10 μM and 30 μM of the free RNA oligos dissolved in RNase free water. The preincubation step was followed by incubation with 20 μL settled beads for 30 min at 4 °C. The water control lysate was recovered and incubated similarly with RNA oligo beads as a pull down of pull down experiment to calculate the depletion factor. The bound proteins were subsequently eluted with 60 μL of 2× NuPAGE LDS sample buffer (Invitrogen,Germany) containing 50 mM DTT. Eluates were alkylated with CAA and in-gel digestion were performed. The resulting peptides were measure using nanoflow LC-MS/MS by directly coupling a nanoLC-Ultra 1D+ (Eksigent) to an Orbitrap Elite mass spectrometer (Thermo Fisher Scientific). Peptides were delivered to a trap column (75 μm × 2 cm, self-packed with Reprosil-Pur C18 ODS-3 5 μm resin, Dr. Maisch, Ammerbuch) at a flow rate of 5 μL/min in solvent A (0.1% formic acid in water). Peptides were separated on an analytical column (75 μm × 40 cm, self-packed with Reprosil-Gold C18, 3 μm resin, Dr. Maisch, Ammerbuch) using a 100 min linear gradient from 4-32% solvent B (0.1% formic acid, 5% DMSO in acetonitrile) in solvent A_1_ (0.1% formic acid, 5% DMSO in water) at a flow rate of 300 nL/min (Hahne *et al*, 2013) Full scans (m/z 360-1,300) were acquired at a resolution of 30,000 in the Orbitrap using an AGC target value of 1e6 and maximum injection time of 100 ms. Tandem mass spectra were generated for up to 15 peptide precursors were selected for fragmentation by higher energy collision-induced dissociation (HCD) using 30% normalized collision energy (NCE) and analyzed in the Orbitrap at a resolution of 7,500 resolution using AGC value of 2e5 and maximum injection time of 100 ms. For peptide and protein identification and label free quantification, the MaxQuant suite of tools version 1.5.3.30 was used. The spectra were searched against the Uniprot human proteome database with carbamidomethyl (C) specified as a fixed modification. Oxidation (M) and Acetylation (Protein N-Term) were considered as variable modifications. Trypsin/P was specified as the proteolytic enzyme with 2 maximum missed cleavages. Label-free quantification (Cox *et al*, 2014) and the match between runs function was enabled. The FDR was set to 1% at both PSM and protein level. Protein intensities were normalized to the respective water control and IC50 and EC50 values were deduced by a four-parameter log-logistic regression using an internal pipeline that utilizes the ‘drc’ package (Ritz *et al*, 2015). A Kd was calculated by multiplying the estimated EC50 with a protein-dependent correction factor (depletion factor) as previously described (Médard *et al*, 2015).

### Data availability

Transcriptome sequencing and quantification data are available at www.ebi.ac.uk/arrayexpress/experiments/E-MTAB-2836/. The raw mass spectrometric data and the MaxQuant result files are available from the Proteomics IDEntifications (PRIDE, http://www.ebi.ac.uk/pride) database (accession number: PXD010153 and PXD010154).

#### Acknowledgements

We thank Stephanie Heinzlmeir for valuable discussions on the RNA-motif competition binding assay. We thank Jun Cheng for help on the RNA half-life estimations. We thank Terence Hwa for discussion on prokaryotic transcriptome and proteome relationship. This work was in part funded by the German Excellence Initiative cluster Center for Integrated Protein Analysis Munich (CIPSM). Dongxue Wang is grateful for a scholarship from the Chinese Research Council. Basak Eraslan has been funded by QBM scholarship. Holger Prokisch has been funded by SOUND research project

## Author contributions

MU, HP, HH, BK, JG conceived and designed the study

DW, MG, AA performed experiments

BE, DW, TW, HH, BK, JG conceived the data analysis

BE, DW, BH, TW, TH, JG analyzed data

BE, DW, MG, HH, BK, JG, TW wrote the manuscript

FP contributed samples and data to the project

## Expanded View Figure legends

**Figure EV1.**
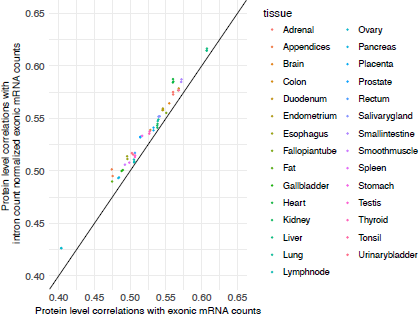
Comparison of spearman correlations between tissue specific protein (iBAQ (log^10^)) and mRNA levels (length normalized exonic read counts (log^10^)) without (x-axis) and with (y-axis) intronic read count normalization.

**Figure EV2.**
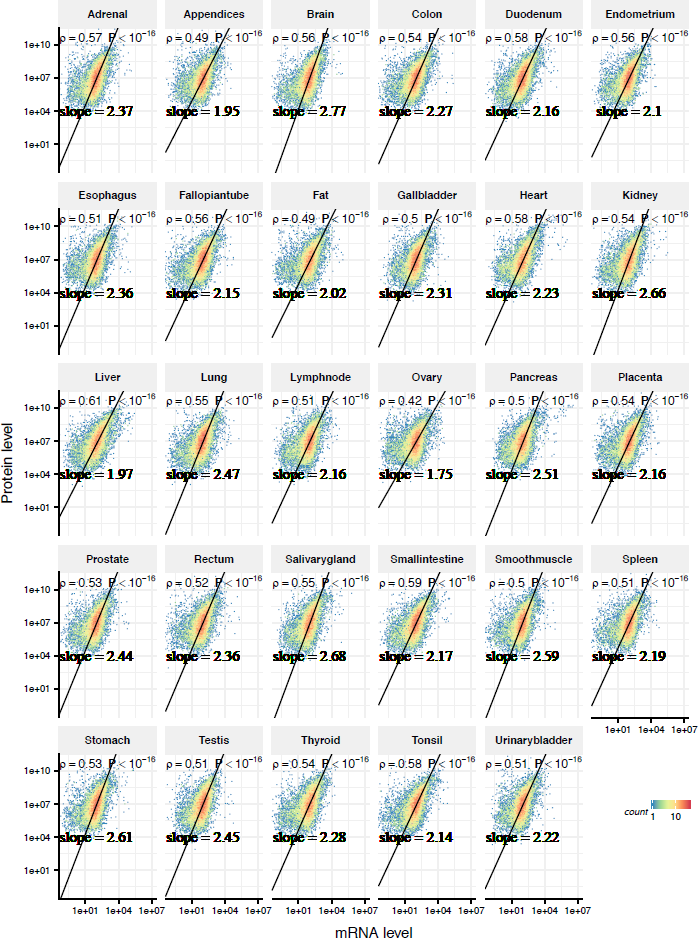
Tissue specific mRNA (x-axis) and protein (y-axis) levels, with spearman correlation coefficients and ranged major axis fit (black line) per tissue.

**Figure EV3.**
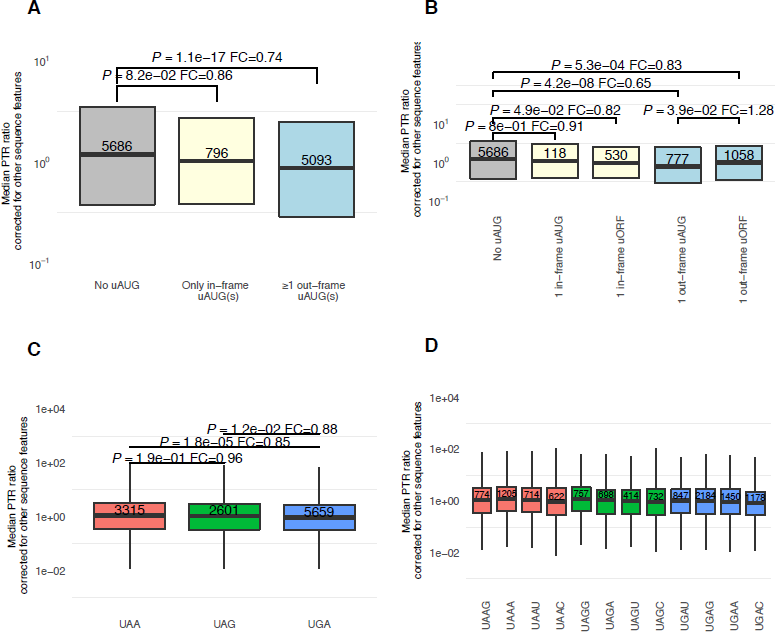
A Transcripts having at least 1 out-frame uAUG has significantly less PTR ratio across 29 tissues (corrected for other sequence elements) compared to transcripts with either no uAUG or only with in-frame uAUG(s) (Wilcox test FC=0.75, P=6.8e-17) B Transcripts with only one out-frame start codon which is not followed by an in frame stop codon in the 5’ UTR (uAUG) is associated with 22% less PTR ratio (corrected for other sequence elements) compared to the transcripts with only 1 out-frame upstream open reading frame (uORF) (Wilcox test FC=0.78, P=3.9e-02). C Transcripts with opal (UGA) canonical stop codon has significantly less median PTR ratio across 29 tissues (corrected for other sequence elements) compared to transcripts ending with ochre (UAA) or amber (UAG). D Transcripts with a cytosine at +1 position has significantly less median PTR ratio across 29 tissues (corrected for other sequence elements) independent of the stop codon type.

**Figure EV4.**
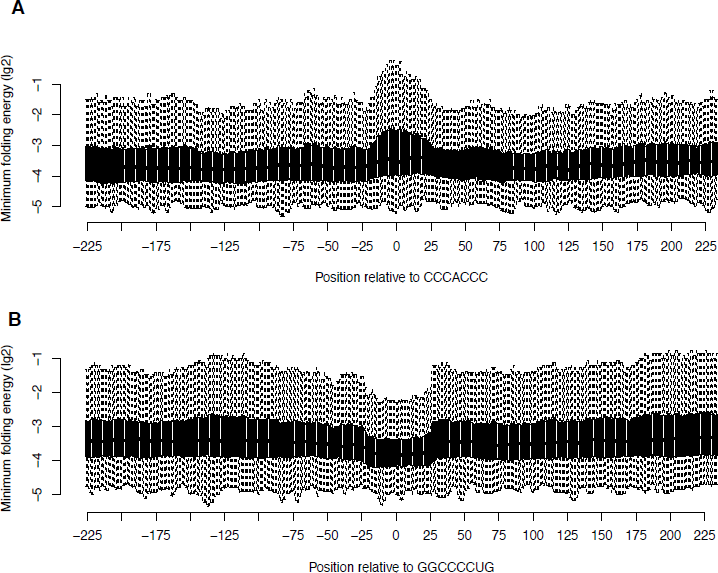
A Negative absolute minimum folding energy (log^2^) distributions of 51 nucleotide sequences centered at the position relative to the CCCACCC sites (x-axis) B Same as A, for GGCCCCUG sites (x-axis)

**Figure EV5.**
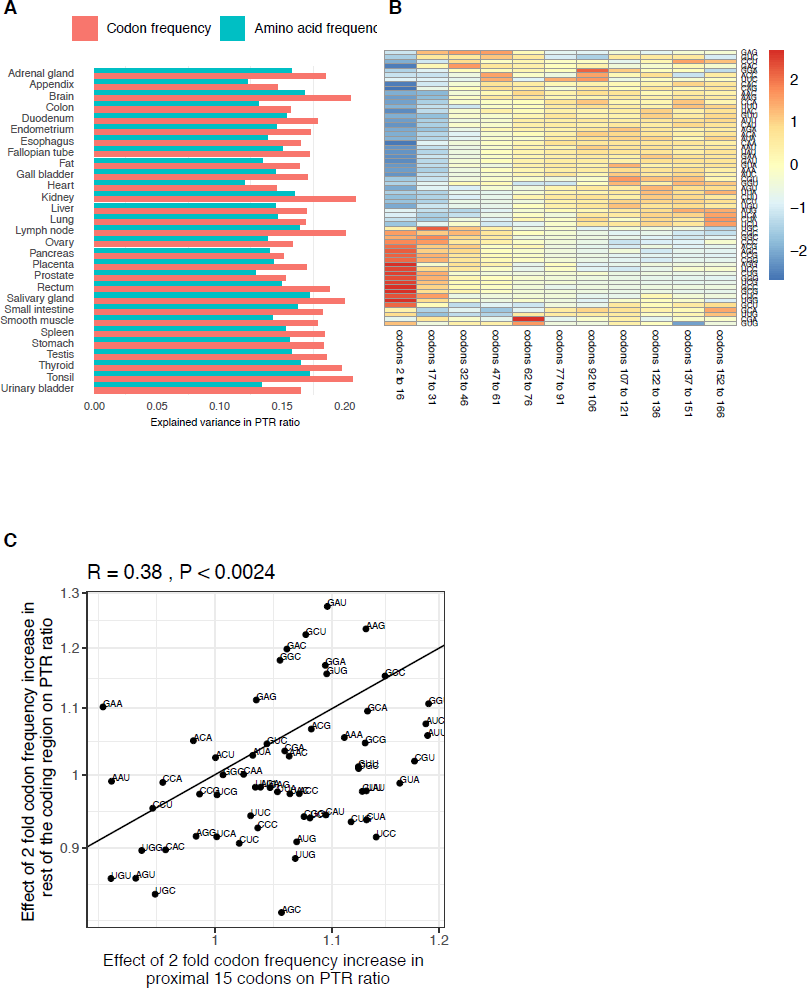
A Explained variance in PTR ratio by log_2_ frequency of 61 codons is higher than log_2_ frequency of 20 amino acids in all 29 tissues. B Codon frequencies relative to average codon frequency (log_2_) per bin of 15 codons (columns). Codon frequency is differentially distributed in the 5’ end of the CDS (from codon 2 to 31) compared to the rest of the coding region. C In spite of the distinctive codon frequency composition of the coding region 5’ end, relative codon effect on the PTR ratio is highly correlating with the codon effects in the rest of the coding region (Spearman’s correlation=0.38, P < 0.0024)

**Figure EV6.**
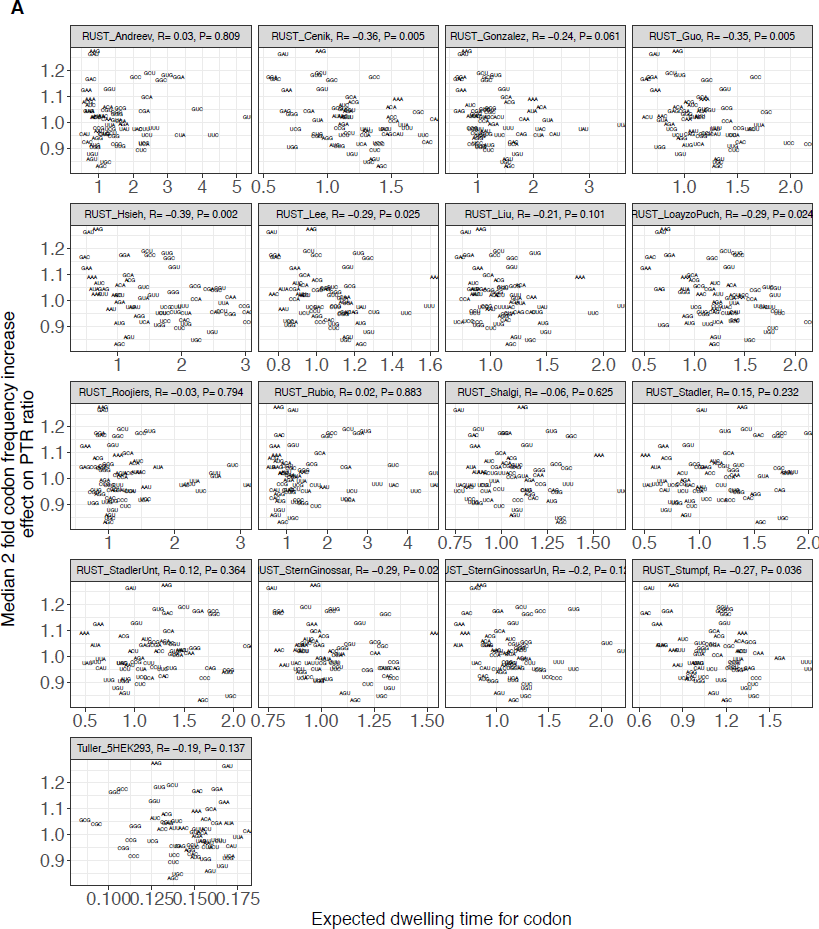
A Comparison of median effect of codon 2 fold frequency increase on PTR ratios across tissues with expected codon dwelling times by RUST (O’Connor *et al*, 2016) based on various ribosome profiling data sets.

**Figure EV7.**
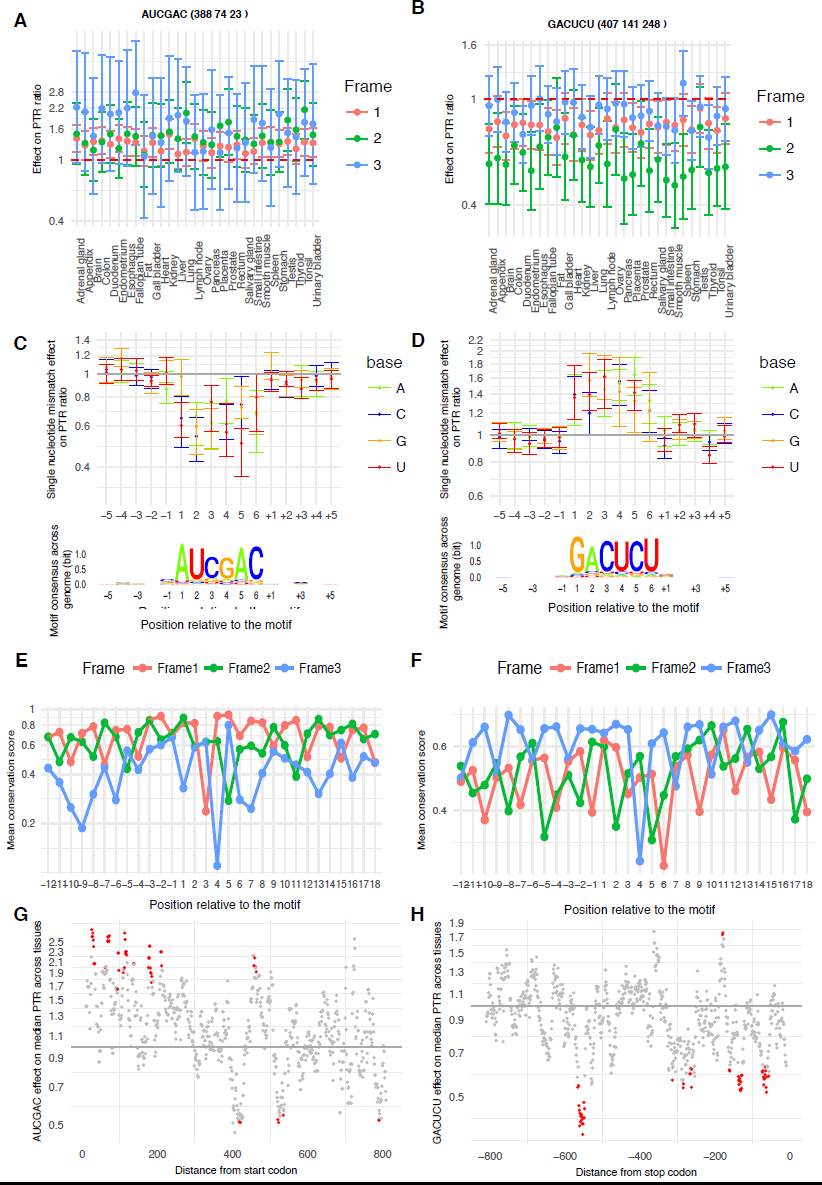
A Frame specific effect estimate (dot) and 95% confidence interval (bar) of the number of perfect match of the N-terminal ([0, + 300] to the canonical start codon) 6-mer AUCGAC on tissue specific PTR ratios. AUCGAC is available in the N-terminal sites of 388,74 and 23 transcripts in frame 1(canonical coding frame), 2 and 3 respectively. B Same as A for C-terminal ([-300, 0] to the canonical stop codon) 6-mer GACUCU. C (Top) Median effect (dot) and across tissue range (bar) of a single mismatch per position relative to AUCGAC on log10 PTR ratio corrected for all other sequence features. (Bottom) Position weight matrix logo of AUCGAC showing information in bits (y-axis) computed across all motif instances in coding region N-terminal window [0, + 300] with respect to the canonical start codon, allowing for at most one mismatch. D Same as C for GACUCU for C-terminal window [-300, 0] with respect to the canonical stop codon. E Average 100-vertebrate PhastCons score (y-axis, Materials and Methods) per position per motif frame relative to the exact AUCGAC match instances in coding region N-terminal end(x-axis) F Same as E for GACUCU for C-terminal window [-300, 0]. G Effect size and effect significance (red FDR<0.1) of aucgac on median ptr ratio across tissues based on its distance to canonical start codon. H Same as G for GACUCU based on its distance to canonical stop codon.

**Figure EV8.**
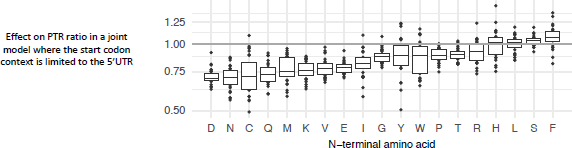
Effects of protein N-terminal residues on PTR ratio compared to Alanine.

**Figure EV9.**
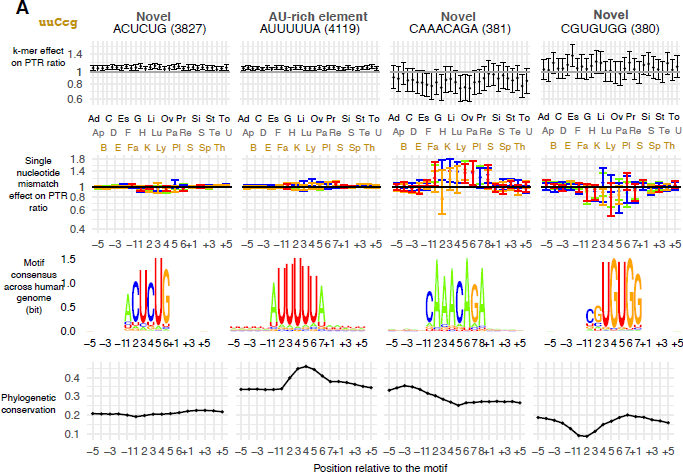
Same as Fig 2E, for 4 3′ UTR motifs.

**Figure EV10.**
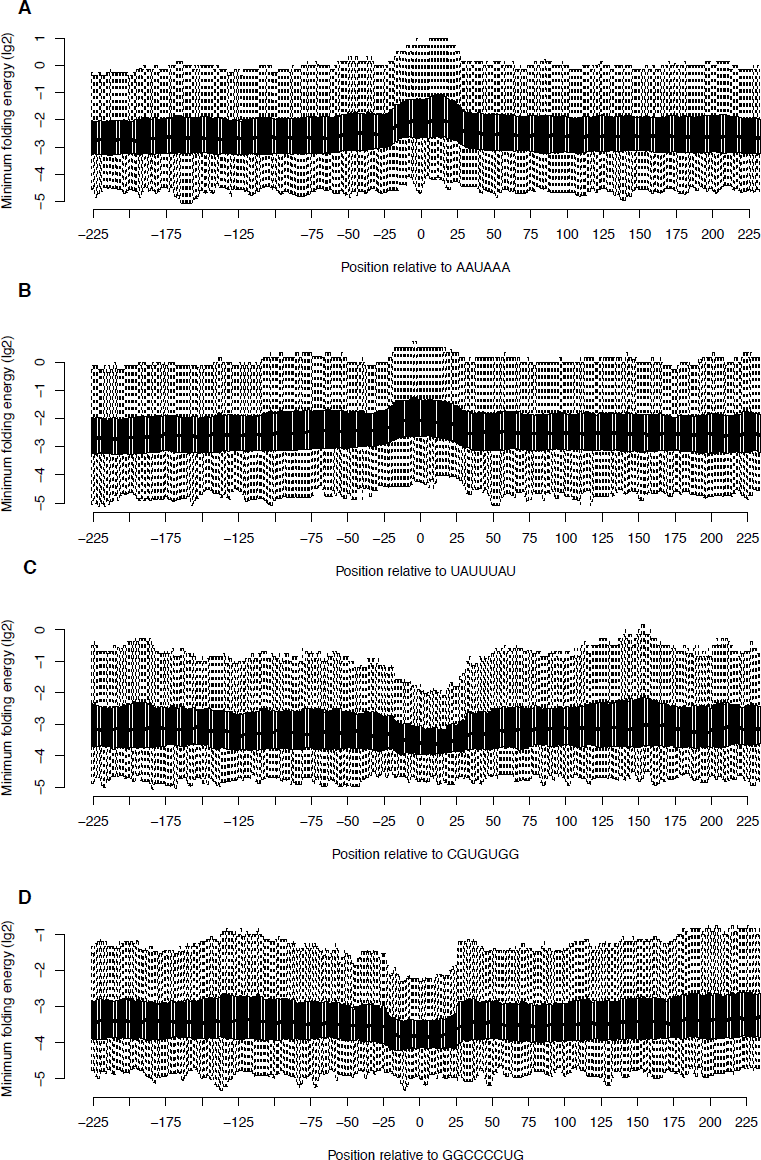
Same as Fig EV4, for 3′ UTR motifs AAUAAA, UAUUUAU, CGUGUGG and GGCCCCUG.

**Figure EV11.**
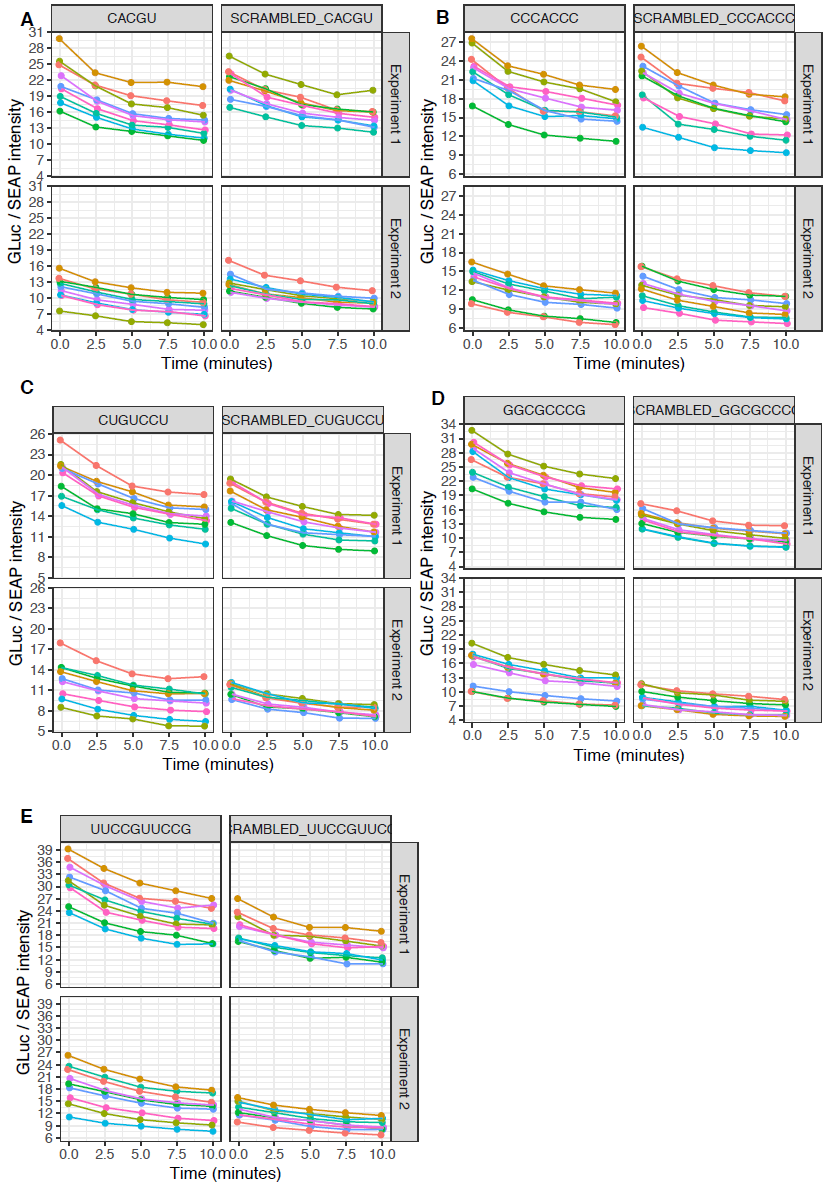
Time course Gluc/SEAP intensity values per 5′ UTR motif and its scrambled version.

**Figure EV12.**
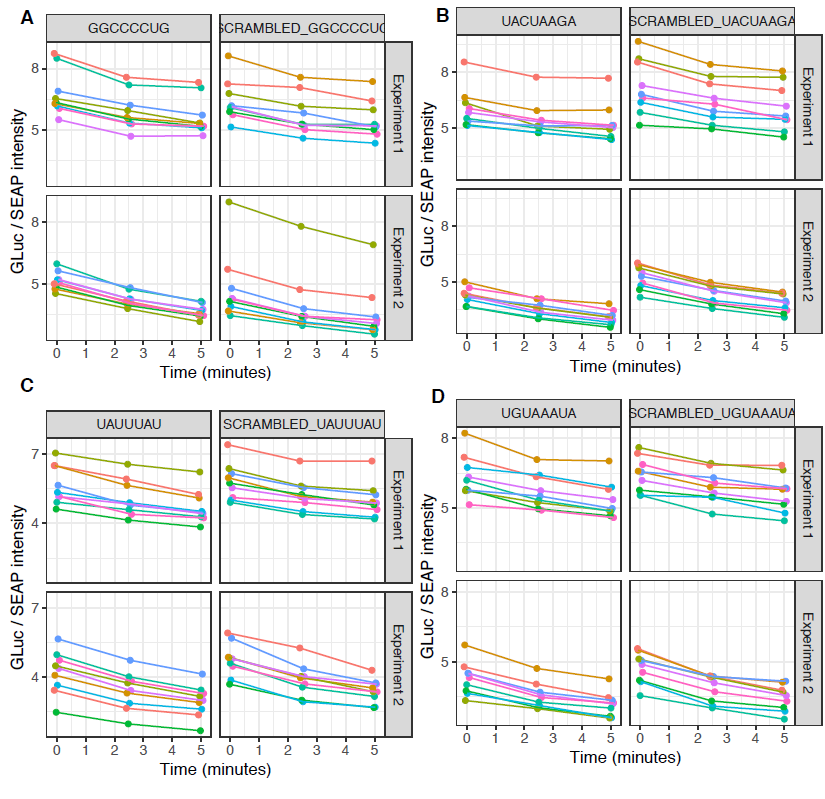
Time course Gluc/SEAP intensity values per 3′ UTR motif and its scrambled version.

